# Macrophages govern antiviral responses in human lung tissues protected from SARS-CoV-2 infection

**DOI:** 10.1101/2021.07.17.452554

**Authors:** Devin J. Kenney, Aoife K. O’Connell, Jacquelyn Turcinovic, Paige Montanaro, Ryan M. Hekman, Tomokazu Tamura, Andrew R. Berneshawi, Thomas R. Cafiero, Salam Al Abdullatif, Benjamin Blum, Stanley I. Goldstein, Brigitte L. Heller, Hans P. Gertje, Esther Bullitt, Alexander J. Trachtenberg, Elizabeth Chavez, Amira Sheikh, Susanna Kurnick, Kyle Grosz, Markus Bosmann, Maria Ericsson, Bertrand R. Huber, Mohsan Saeed, Alejandro B. Balazs, Kevin P. Francis, Alexander Klose, Neal Paragas, Joshua D. Campbell, John H. Connor, Andrew Emili, Nicholas A. Crossland, Alexander Ploss, Florian Douam

**Affiliations:** Department of Microbiology, Boston University School of Medicine, Boston, MA, USA; National Emerging Infectious Diseases Laboratories, Boston University, Boston, MA, USA; Bioinformatics Program, Boston University, Boston, MA, USA; Department of Pathology and Laboratory Medicine, Boston University School of Medicine, Boston, MA, USA; Center for Network Systems Biology, Boston University, Boston, MA, USA; Department of Biochemistry, Boston University School of Medicine, Boston, MA, USA; Department of Molecular Biology, Princeton University, Princeton, NJ, USA; Single Cell RNA Sequencing Core, Boston University, Boston, MA, USA; Department of Medicine, Boston University School of Medicine, Boston, MA, USA; Department of Physiology and Biophysics, Boston University School of Medicine, Boston, MA, USA; Animal Science Center, Boston University, Boston, MA, USA; Center for Thrombosis and Hemostasis, University Medical Center of the Johannes Gutenberg-University, Mainz, 55131, Germany; Electron Microscopy Core Facility, Harvard Medical School, Boston, MA, USA; Department of Neurology, Boston University School of Medicine, Boston, MA, USA; Ragon Institute of MGH, MIT and Harvard, Cambridge, MA, USA; PerkinElmer, Hopkinton, MA, USA; In Vivo Analytics, Inc., New York, NY, USA; Department of Radiology Imaging Research Lab, University of Washington, Seattle, WA, USA; Department of Biology, Boston University School of Medicine, Boston, MA, USA

## Abstract

The majority of SARS-CoV-2 infections among healthy individuals result in asymptomatic to mild disease. However, the immunological mechanisms defining effective lung tissue protection from SARS-CoV-2 infection remain elusive. Unlike mice solely engrafted with human fetal lung xenograft (fLX), mice co-engrafted with fLX and a myeloid-enhanced human immune system (HNFL mice) are protected against SARS-CoV-2 infection, severe inflammation, and histopathology. Effective control of viral infection in HNFL mice associated with significant macrophage infiltration, and the induction of a potent macrophage-mediated interferon response. The pronounced upregulation of the USP18-ISG15 axis (a negative regulator of IFN responses), by macrophages was unique to HNFL mice and represented a prominent correlate of reduced inflammation and histopathology. Altogether, our work shed light on unique cellular and molecular correlates of lung tissue protection during SARS-CoV-2 infection, and underscores macrophage IFN responses as prime targets for developing immunotherapies against coronavirus respiratory diseases.

**HIGHLIGHTS:**

- Mice engrafted with human fetal lung xenografts (fLX-mice) are highly susceptible to SARS-CoV-2.
- Co-engraftment with a human myeloid-enriched immune system protected fLX-mice against infection.
- Tissue protection was defined by a potent and well-balanced antiviral response mediated by infiltrating macrophages.
- Protective IFN response was dominated by the upregulation of the USP18-ISG15 axis.

## INTRODUCTION

In December 2019, the emergence of coronavirus disease 19 (COVID-19), a new viral respiratory disease (Zhu et al., 2020), created a public health emergency and economic disruption at an unprecedented scale. As of July 6^th^, 2021, the etiologic agent of COVID-19, severe acute respiratory syndrome coronavirus 2 (SARS-CoV-2), a β-Coronavirus from the *Coronaviridae* family, has infected more than 183 million individuals and caused nearly 4 million deaths (WHO COVID-19 Weekly Epidemiological Update; July 6^th^, 2021). The COVID-19 pandemic illustrates the necessity of increasing our knowledge of *Coronaviridae* to better anticipate potential future coronavirus pandemics.

Human immune correlates of severe COVID-19 disease have been intensively investigated since the onset of the pandemic. A consensual model suggests that severe COVID-19 is associated with profound dysregulation of host immune responses, which exacerbates tissue damage and inflammation (Vabret et al., 2020). These dysregulations include disordered T-cell effector function and lower mobilization (Wauters et al., 2021), lung enrichment in hyperinflammatory monocytes, interstitial macrophages, and neutrophils (Rendeiro et al., 2021; Wauters et al., 2021), low interferon (IFN) responses (Combes et al., 2021; Hadjadj et al., 2020), increased production of autoantibodies (Bastard et al., 2020; Combes et al., 2021), extensive complement activation (Ma et al., 2021), delayed neutralizing antibody responses (Lucas et al., 2021) or/and dysfunctional tissue repair mechanisms (Delorey et al., 2021).

However, in young, healthy individuals, most SARS-CoV-2 infections result in asymptomatic to mild disease (Nikolai et al., 2020; Wu and McGoogan, 2020), suggesting the induction of protective immune responses against SARS-CoV-2. While understanding the molecular basis of such protective responses could reveal major avenues for the development of effective immunotherapies against COVID-19, conducting such studies has been hampered by several roadblocks. Indeed, studies on patients with asymptomatic to mild infections carry ethical limitations that restrict immunological sampling to the peripheral blood, bronchioalveolar fluid (BALF), and nasopharynx, and these sampling strategies also contain critical constraints. For example, peripheral blood only represents a subset of the immune responses occurring in the respiratory tract upon infection, and BALF/nasopharynx swabs do not provide access to histopathological and spatial information. Additionally, the interpretation of these studies is confounded by several factors, including non-synchronized collection of tissues post-infection, inter-individual variability and co-morbidities, differences in viral inoculum and route of infection, and exposure to different viral strains.

The inherent ethical limitations associated with human studies highlights animal models of infection as a valuable alternative to better understand the molecular mechanisms driving COVID-19 disease and protection. Animal models not only provide increased control of experimental settings, they also allow us to go beyond purely descriptive studies and to experimentally challenge hypotheses. Non-human primates (NHPs), hamsters, and ferrets are naturally susceptible to SARS-CoV-2 infection and have been instrumental in evaluating the therapeutic potential and prophylactic efficacy of many antiviral countermeasures (Kim et al., 2020; Sia et al., 2020; Singh et al., 2021; Tostanoski et al., 2020; Vogel et al., 2021). However, these models have also demonstrated limitations for in-depth analysis of coronavirus pathogenesis and immunity. COVID-19 mostly causes only subclinical disease in NHPs (Singh et al., 2021), and the scarcity of proper reagents and the cost-prohibitive aspects of such studies collectively limit the use of these animals for deep and comprehensive immunological investigations.

In parallel, the large evolutionary divergence between humans and hamsters or ferrets, manifesting in vastly different immune responses and lung environments, makes studies in these animals unlikely to recapitulate the complex array of human-virus interactions that drive immune responses to SARS-CoV-2. Similar limitations are present in murine models, as reported in mice transgenically expressing or harboring a knock-in for the human orthologs of angiotensin-converting enzyme 2 (ACE2), the cell entry receptor of SARS-CoV-1 and -2 (Carossino et al., 2021; Jiang et al., 2020; Winkler et al., 2020; Zheng et al., 2021). Additionally, there are concerns about the hyper-susceptibility to infection displayed by some of these models (Carossino et al., 2021).

Mice harboring human tissue xenografts have been used to investigate a variety of infectious diseases over the past three decades (Douam and Ploss, 2018). Recently, mice engrafted with human fetal lung xenografts (fLX) were successfully infected with several coronaviruses including SARS-CoV-2 (Wahl et al., 2019; Wahl et al., 2021). However, the potential of these models to dissect human hematopoietic responses to SARS-CoV-2 has not been explored. Here, we harnessed fLX-engrafted mice, engrafted or not with human hematopoietic components, to identify immunological correlates of lung tissue protection (i.e. protection from viral infection, and from lung tissue damage) from SARS-CoV-2 infection. Immunodeficient NOD-*Rag1^-/-^IL2Rγ^NUL^* (NRG) mice, or NRG-*Flk2-/-*Flt3LG mice (Douam et al., 2018) engrafted with components of a human immune system, were engrafted with pairs of fLX, yielding NRG-L and HIS-NRGF/Flt3LG-L (referred to as HNFL) mice, respectively. Viral inoculation of fLX in NRG-L mice resulted in robust type-I IFN responses and prolonged inflammation, which associated with persistent infection and severe histopathology. In contrast, HNFL mice were able to control SARS-CoV-2 infection without exhibiting severe inflammation and histopathology. Protection in HNFL mice was associated with significant macrophage infiltration and by the induction of a strong macrophage-mediated IFN response, which was dominantly characterized by the upregulation of the USP18-ISG15 axis – a negative regulator of type I IFN responses and ISGylation (Basters et al., 2018). The USP18-ISG15 upregulation was unique to HNFL mice and was a prominent correlate of limited histopathology and inflammation. Altogether, our work sheds light on the crucial role of macrophage infiltration and macrophage-mediated immunoregulations in driving effective and balanced antiviral and anti-inflammatory responses during SARS-CoV-2 infection, and provides new evidence that dysregulation of macrophage responses can contribute to severe COVID-19 disease. Our work highlights macrophage-mediated IFN responses as promising targets for the development of immunotherapies against COVID-19 and other severe respiratory viral diseases.

## RESULTS

### Human fetal lung xenografts display heterogeneously mature epithelia and resident hematopoietic lineages upon engraftment in NRG mice

Ten to fifteen-week-old male and female NOD-*Rag1^-/-^IL2Rγ^NULL^* (NRG) mice were surgically implanted subcutaneously with two pieces of fLX, one on each side of the animal’s thoracic body wall (**Figure S1A**). Following engraftment, all animals were healthy and displayed palpable and macroscopically noticeable fLX on both sides of their body (**Figure S1A,B**). Macroscopic and histological analysis illustrated interstitial infiltration of fLX by murine blood vessels with retention of human vasculature (**Figure S1C-E**). Cells appeared histologically normal without any evidence of degeneration and/or necrosis that would be suggestive of tissue ischemia or hypoxia (**Figure S1F-M**). The fLX were characterized by sporadic regions of conducing zone differentiation evidenced by the presence of bronchioles and hyaline cartilage, as well as more abundant and heterogenous stages of respiratory zone development, which included combinations of pseudo glandular, canalicular, and saccular phenotypes (**Figure S1F-M**). Different stages of fetal lung maturation were associated with differential expression of ACE2, prosurfactant protein C (SFTPC), an alveolar type 2 (AT2) pneumocyte differentiation marker, and human specific CD31 (blood vessel marker) (**Figure S1F-M**). Regions with a more immature phenotype (pseudo glandular) were associated with the highest immunoreactivity for these three antigens, while more mature stages of fetal development (canalicular and saccular) had decreased expression (**Figure S1F-M**). In regions of respiratory zone differentiation, ACE2 expression was primarily restricted to cells concurrently expressing SFTPC, consistent with AT2 pneumocyte origin. ACE2 was also observed along the apical surface of the bronchiole epithelium, which lacked SFTPC expression. To further characterize the cellular composition of the fLXs, we performed single-cell (sc) RNA sequencing. The human cellular compartment represented 67.14% of the total fLX (mouse+human cells) and was mainly composed of cells from the epithelial lineage (50.01% of the human compartment; **Figure 1A-D**), specifically AT1, AT2, club cells, and ciliated cells (**Figure 1A,C; Supplemental Item 1**). Strikingly, we also detected a human resident hematopoietic compartment in fLX (13.26%) which was composed of a myeloblast progenitor cluster (c-kit^+^ SRGN^+^ CD14^+^ HPGD^+^; 3.13%) and a lymphoid cluster with cytotoxic functions (CD8^+^ T cells and/or NK cells; 10.13%). These findings were confirmed by flow cytometry (**Figure S2A-D**). As expected, the mouse compartment in fLX was mainly composed of cells from the myeloid lineages (82.26% rep.) (**Figure 1B,D**). Altogether, our findings unravel the complex cellular composition of fLX, which extends beyond the human pulmonary epithelial compartment.

**Figure 1.**
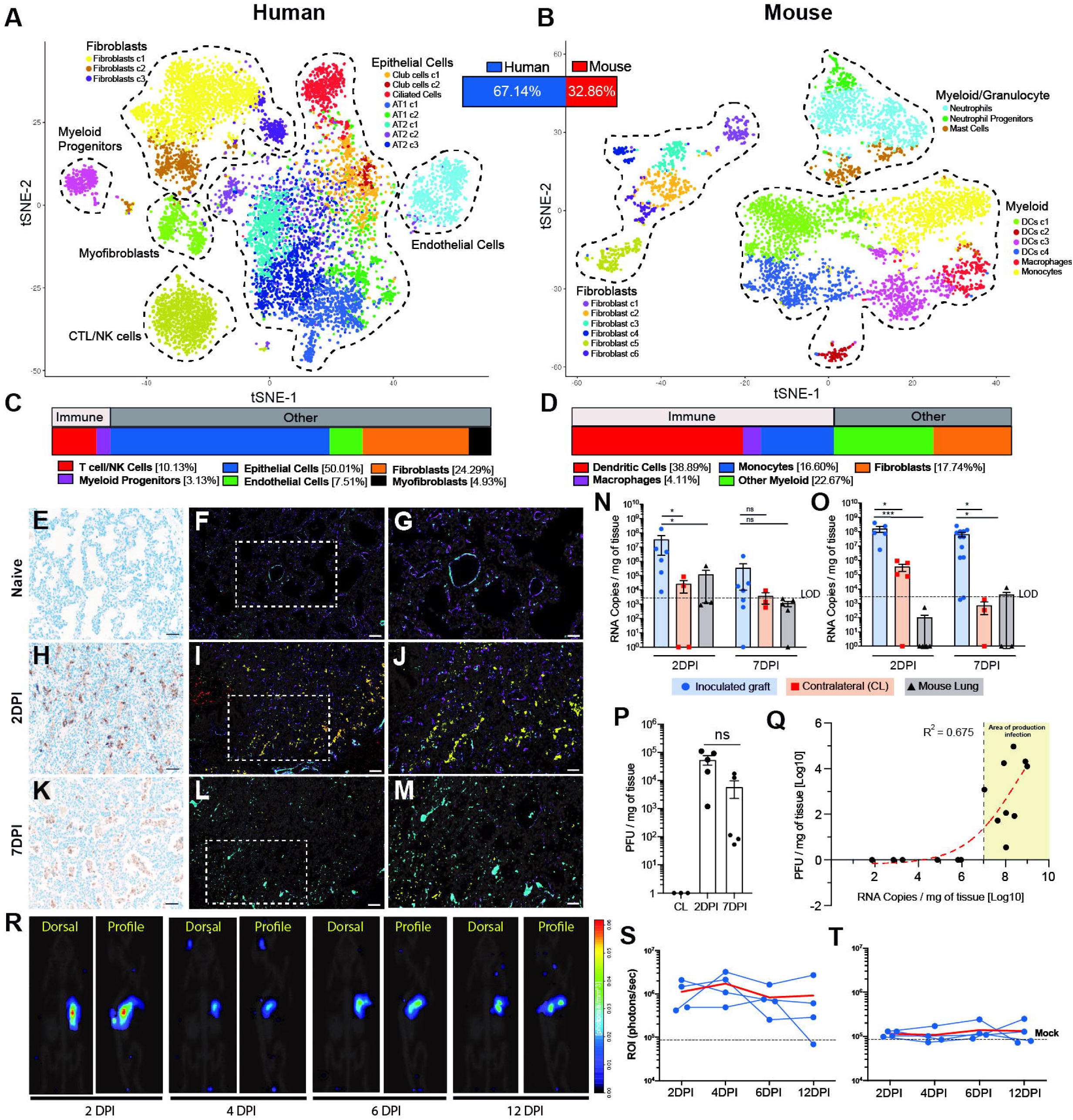
NRG-L mice are susceptible to SARS-CoV-2 infection. See also Figure S1 and S2, and Supplemental Items 1 and 2. **A-D.** t-SNE plot of the human (A: 3 fLX, 9,968 cells) and mouse (B: 3 fLX, 5,091 cells) compartment in fLX of NRG-L mice. Relative representation of each compartment within fLX is indicated as a percentage between the two t-SNE plots. Cell subset frequencies within each compartment (C: human; D: mouse) are shown below the respective t-SNE plot. **E-M.** Representative SARS-CoV-2 N IHC (E,H,K) and 5-color IHC (F,G,I,J,L,M: yellow: SARS-CoV-2 Spike; Magenta, human CD31; Cyan, human CD61; Red, human CD68; Grey, DAPI) on non-inoculated (E,F,G: Naïve fLX), or inoculated fLX tissue section (H,I,J: 2DPI; K,L,M: 7DPI) from NRG-L mice using 10^6^ PFU of SARS-CoV-2. G, J, M are a 2x magnification of the inset located in F, I, L respectively. F,I,L: 100x, scale bar=200µm; E,H,K: 200x, scale bar=100µm; G,J,M: 400x, scale bar=50 µm. **N-O.** SARS-CoV-2 viral RNA quantification by RT-qPCR in inoculated graft (blue), contralateral non-inoculated graft (red) and in mouse lung (grey) using 10^4^ (N) or 10^6^ (O) SARS-CoV-2 PFU (n=4-12 fLX). Limit of detection (LOD) is shown as a dotted line and is equivalent to RNA copies/mg tissues in naïve fLX (n=7). Mean±SEM, Kruskal-Wallis test; *p≤0.05, \*\*\**p*≤0.001. **P.** Quantification of infectious SARS-CoV-2 particles by plaque assay in non-inoculated contralateral (CL, 7DPI) or inoculated fLX (2DPI and 7DPI) with 10^6^ PFU of SARS-CoV-2. n=3-5 fLX. Mean±SEM, Kruskal-Wallis test; ns, non-significant. **Q.** Non-linear regression (Sigmoidal 4PL) between viral RNA copies and infectivity per mg of tissue in inoculated (n=10; 2DPI and 7DPI; 10^6^ PFU), fLX contralateral fLX (n=3) and in naïve fLX mice (n=2). The yellow area associates with productive infection and starts at the productive infection threshold (PIT) established at 10^7^ viral RNA copies/mg tissue. n=15 fLX. **R.** Representative three-dimensional dorsal and profile view of a single NRG-L mouse following inoculation of the right fLX with a SARS-CoV-2 NanoLuc virus (10^6^ PFU), over 12 days of infection. NanoLuc bioluminescent signal was detected and quantified over time using the InVivoPLOT (InVivoAx) system and an IVIS Spectrum (PerkinElmer) optical imaging instrument. **S-T**. Regionalized quantification of NanoLuc expression in inoculated right fLX (S; n=4) and in non-inoculated contralateral fLX (T: n=4). Quantification was performed over a 12-day course of infection. Mean signal from naïve fLX (n=3) was used to determine assay baseline (mock). Red line represents the mean signal over time.

### Persistent SARS-CoV-2 infection in NRG-L mice following fLX inoculation

Ten to fifteen weeks post-engraftment, the NRG-L mice were inoculated with a SARS-CoV-2 clinical isolate (USA-WA1/2020 isolate) at a viral dose of either 10^4^ or 10^6^ plaque forming units (PFU). To evaluate the possibility of fLX-to-fLX viral transfer through the peripheral blood, some animals were inoculated only in a single fLX. Non-inoculated fLX from non-infected and infected animals are hereafter referred to as “naïve” and “contralateral” fLX respectively. Over the course of infection, NRG-L mice did not display any signs of clinical disease, maintained their weight, and had no significant alternations in body temperature (**Figure S2E-F**). SARS-CoV-2 inoculation resulted in the development of gross abnormalities in fLX (**Figure S2H**), which contrasted with a white homogenous appearance of contralateral fLX.

Immunoreactivity to SARS-CoV-2 nucleocapsid (N) in inoculated fLX was observed in a dose dependent manner, with more abundant viral antigen observed in fLX inoculated with the 10^6^ dose as compared to 10^4^ dose (**Figure 1E,H,K**; **Figure S2I-O**). When challenged with 10^6^ PFU, viral antigen was concentrated in pneumocytes and bronchiole epithelium at 2 days post inoculation (DPI) (**Figure 1H; Figure S2N,P**), although at 7DPI it was mainly found in necrotic debris within the airspaces (**Figure 1K; Figure S2O**). Infection appeared to be cleared at 7DPI for the 10^4^ PFU dose (**Figure S2K,L**). These findings were supported by 5-color fluorescent imaging (DAPI, SARS-CoV-2 Spike, CD31, CD61, CD68) (**Figure 1F,G,I,J,L,M**), which also showed an increase of platelet rich thrombi in infected fLX at 7DPI, a feature consistent with previous studies in human and non-human primates (Aid et al., 2020; Mackman et al., 2020).

RT-qPCR quantification of viral RNA (E gene) was consistent with immunohistochemistry (IHC) findings (**Figure 1N-O**). Although RT-qPCR suggested that low levels of viral RNA were present in contralateral fLX, SARS-CoV-2 N immunoreactivity was never observed in contralateral fLX (**Figure S2Q**). Quantification of infectious viral particles at 2DPI and 7DPI supported evidence of productive infection in fLX, with detection of infectious viral particles at both time points in infected fLX. No infectious viral particles were detected in contralateral fLX (**Figure 1P**) suggesting that limited amount of viral RNA but not infectious viral particles may transfer between fLX through the blood. However, no viral RNA was detected in the peripheral blood at any time point (**Figure S2R**) further strengthening evidence that the amount of circulating viral RNA transferring between fLX is probably very low.

We found a positive non-linear regression between viral load and PFU in infected fLX (R^2^=0.675) (**Figure 1Q**). Using these data, we identified that viral loads in excess of a threshold of 10^7^ RNA copies/mg tissue (i.e. productive infection threshold or PIT) were indicative of productive infection.

To further confirm evidence of productive infection in NRG-L fLX, we performed planar and 3D *in vivo* bioluminescence imaging on NRG-L mice inoculated with a recombinant SARS-CoV-2 expressing NanoLuc luciferase (Xie et al., 2020). The NanoLuc bioluminescent signal was readily detectable in inoculated fLX mice as early as 2DPI (**Figure 1R,S; Figure S2S-T; Supplemental Item 2**) but not contralateral grafts (**Figure 1R-T; Figure S2S,T**), which was consistent with our findings above. Most importantly, SARS-CoV-2 replication was maintained in single animals for up to 12DPI. Altogether, our findings demonstrate that fLX support productive and persistent SARS-CoV-2 infection.

### Persistent infection in NRG-L mice associates with extensive histopathological manifestations

To capture the heterogeneity and severity of histologic phenotypes observed in SARS-CoV-2 infected fLX, we developed a semi-quantitative ordinal scoring (**see methods**). The mean cumulative histologic score in animals inoculated with 10^6^ PFU was 1.76-fold higher at 2 DPI and 2.22-fold higher at 7 DPI compared to those inoculated with 10^4^ PFU, indicating a positive correlation with the viral load and SARS-CoV-2 N positivity. (**Figure 2A; Figure S3A**). Therefore, we elected to pursue all subsequent analysis of SARS-CoV-2 infected fLX using the 10^6^ dose, which when compared to naïve fLX, resulted in an increasing histopathological score over time (**Figure 2B; Figure S3B**). Notably, neutrophil influx, intra-airspace necrosis, capillary fibrin thrombi, and presence of syncytial cells were the most significant independent observations that contributed to the increased cumulative score. Of note, tissue integrity and architecture between naïve and contralateral fLX was similar. Therefore, this histopathological analysis, both naïve and fLX were pooled together to define the histopathological baseline **(Figure 2B**, **Figure S3B**).

**Figure 2.**
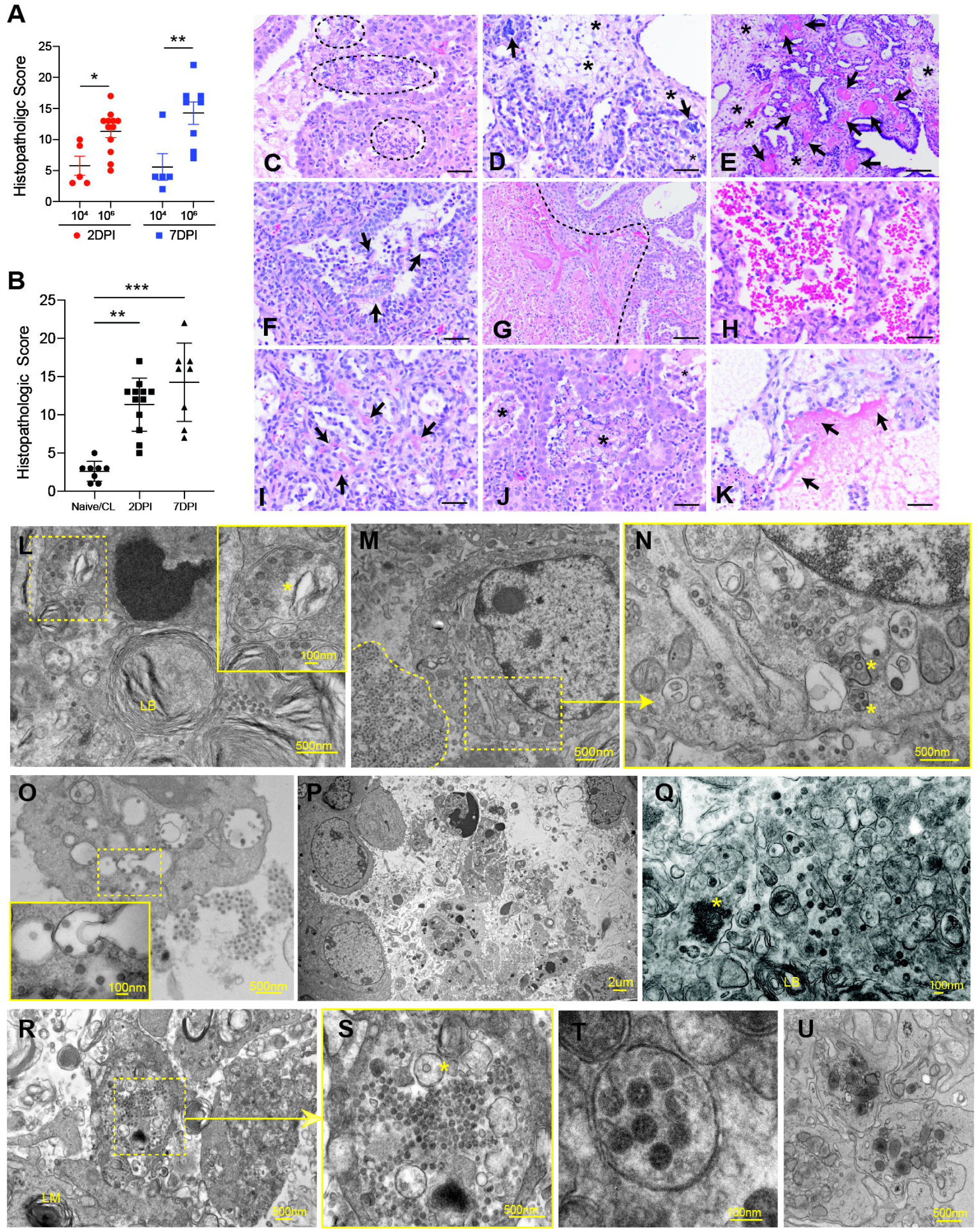
NRG-L inoculation with SARS-CoV-2 results in severe histopathology. See also Figure S3 and S4. **A.** Cumulative histopathologic score of fLX inoculated with 10^4^ or 10^6^ PFU of SARS-CoV-2 at 2DPI and 7DPI. n=5-10 fLX. Mean±SEM, two-way ANOVA; \**p*≤0.05 \*\**p*≤0.01. **B.** Cumulative histopathologic score of fLX inoculated with 10^6^ PFU of SARS-CoV-2 at 2DPI and 7DPI versus naïve/Contralateral (CL) fLX. n=8-12 fLX. Mean±SEM, Kruskal-Wallis test; \*\**p*≤0.01, \*\*\**p*≤0.001. **C-K.** Representative histopathologic phenotypes in fLX inoculated with 10^6^ PFU of SARS-CoV-2 in NRG-L mice. (C) Neutrophil accumulation within airspaces (within black hash lines). (D) Syncytial epithelial cells (black arrows) and interstitial edema (asterisks). (E) Fibrin thrombi occluding intermediate sized blood vessels (black arrows), with interstitial edema (asterisks). (F) Denuded epithelium (black arrows). (G) Coagulative necrosis (left of hash line), adjacent to viable fLX (right of hash line). (H) Intra-airspaces hemorrhage. (I) Fibrin thrombi occluding interstitial capillaries (black arrows). (J) Accumulation of necrotic debris within airspaces (asterisks). (K) Hyaline membrane (black arrows). Hematoxylin and Eosin (H&E) staining, 400x, scale bar=50um. **L-U.** Ultrastructural analysis of inoculated fLX with 10^6^ PFU of SARS-CoV-2. (L) SARS-CoV-2 viral particles in double membrane-bound vesicle (DMV, asterisk) in AT2 pneumocytes (lamellar body, LB) at 2DPI. Top right corner is a magnification of the top left inset (dotted box). (M) Infected pneumocyte with high concentration of viral particles around the peripheral extracellular area (left of hash line) at 2DPI. (N) Magnification (3.75X) of the inset from M with event of viral particle maturation in DMV. (O) Virions free within the airspace phagocytized by a neighboring macrophage with formation of multivesicular bodies containing virions at 2DPI. Bottom left picture is a magnification of the central inset (dotted box). (P) Airspace filled with necrotic cellular debris including lamellar bodies, denuded AT2 pneumocytes undergoing apoptosis characterized by condensation of chromatin and pyknotic nuclei at 7DPI. (Q). Virus particles at variable stages of maturation within the cytoplasm of a type II pneumocyte at 7DPI as indicated by presence of lamellar bodies (LB). Virus particles are both free and in DMV, and electron dense viral replication centers are indicated with an asterisk. (R-S) Airspace filled with necrotic cellular debris at 7DPI including lamellar bodies and anuclear cellular fragments of denuded AT2 pneumocytes containing a DMV (asterisk) and virus particles at varying stages of maturation. S is a magnification (3.75X) of the central inset (dotted box) in R. (T) Cluster of mature virus particles with radiating spikes and aggregates of nucleocapsid protein contained within a DMV at 7DPI. (U) Blood vessel occluded by an aggregate of platelets at 7DPI. Scale bar dimensions are indicated for all pictures.

Histopathologic findings were observed in all SARS-CoV-2 inoculated fLX, ranging from mild focal to severe generalized disease (**Figure 2C-K**). Examination of serial sections confirmed that histologic lesions predominated in areas with SARS-CoV-2 N immunoreactivity. At 2DPI, affected terminal airspaces were infiltrated by neutrophils (**Figure 2C**), with variable amounts of edema (**Figure 2D, E**), denuded epithelial cells (**Figure 2F**), coagulative necrosis (**Figure 2I**) and hemorrhage (**Figure 2J**). In areas of severe neutrophilic inflammation, increased numbers of mitotic figures were observed in pneumocytes supportive of active regeneration (**Figure 2C**), with concurrent pneumocyte degeneration represented by cytoplasmic swelling and vacuolation. Capillaries and intermediate-sized arterioles and arteries were multifocally occluded by fibrin thrombi (**Figure 2E, I**). Similar features to those were observed at 7DPI, but airspaces were more frequently filled with abundant necrotic cellular debris (**Figure 2J**), and in one fLX, a distinctive hyaline membrane lining pneumocytes could also be observed (**Figure 2K**).

Transgenic C57BL/6 mice expressing human ACE2 under the control of the cytokeratine-18 promoter (K18-hACE2) have been reported to be highly susceptible to SARS-CoV-2 (Winkler et al., 2020). Inoculation of K18-hACE2 mice with 10^6^ PFU of SARS-CoV-2 using the same viral stock resulted in multifocal interstitial pneumonia and abundant SARS-CoV-2 S antigen within AT1 and AT2 cells at 2 DPI and increased interstitial histiocyte and lymphocyte infiltrates at 7DPI (**Figure S3C-N**). Although we have recently confirmed that infection with the same viral stock of SARS-CoV-2 was lethal in this model (Carossino et al., 2021), we were unable to observe in this model the diversity and severity of lung histopathological features observed in inoculated fLX, including syncytia, necrosis, microthrombi, and hemorrhage. Taken together, our findings indicate that fLX inoculated with SARS-CoV-2 more faithfully recapitulate several features of diffuse alveolar damage (DAD) as described in cases of severe COVID-19 disease.

Ultrastructural analysis of fLX infected with 10^6^ PFU supported our virological and histopathological findings. Virus particles were observed within AT2 pneumocytes at various stages of maturation and were often confined to double membrane bound vesicles (DMVs), with morphology and particle size consistent with previously described coronaviruses (range 80-130nm in diameter) (Laue et al., 2021) (**Figure 2L-N**). Potential single-particle budding events (**Figure S4A**) and viral particle phagocytosis by macrophages (**Figure 2O**) were observed. At 7DPI, airspaces were filled with abundant necrotic cellular debris including lamellar bodies, erythrocytes, neutrophils and denuded AT2 pneumocytes (**Figure 2P; Figure S4B-D)**, which were occasionally undergoing apoptosis as indicated by the presence of pyknotic nuclei (**Figure S4C**). DMV-containing viral particles and electron-dense viral replication centers were still observed at 7DPI, suggesting persistence of active viral replication (**Figure 2Q-T, Figure S4E**). Faint Spike protein coronal surface projections were sometimes visible within DMVs (**Figure 2T**). Blood vessels also contained aggregates of platelets (**Figure 2U**) with several small to intermediate-sized arteries occasionally occluded by fibrin thrombi (**Figure S4F**). Altogether, our data demonstrate that SARS-CoV-2 infection of NRG-L mice is associated with the development of cellular and histopathological features that resemble those observed in the lungs of severe COVID-19 patients.

### Type-I IFN response and persistent inflammation in infected fLX of NRG-L

COVID-19 has been associated with prolonged inflammation (Blanco-Melo et al., 2020; Rendeiro et al., 2021; Wauters et al., 2021). To further establish the relevance of fLX to recapitulate features of SARS-CoV-2 in the human lung, we examined the transcriptomic and proteomic signatures associated with persistent infection and histopathology. mRNA sequencing yielded mostly human mRNAs; samples with less than 15 million human-aligning reads and/or less than 75% human reads were excluded from further analysis (n=3 out of 19) (**Figure S5A**). As expected, viral RNA was detectable in all inoculated fLX but was more prominently detected in 2DPI samples (**Figure S5B**). Differential expression analyses of human mRNA levels in naive versus 2DPI, 7DPI, and contralateral samples showed evidence of induction of antiviral host responses (**Figure 3A-D; Figure S5C-D; Supplemental Item 3**). A strong type I IFN response was observed at 2DPI, which included transcripts for IFNB1 and L1 as potential drivers of both local and systemic response (**Figure 3A,D; Figure S5C-E**). Chemokines previously reported to be upregulated in severe COVID-19 patients, such as CXCL9, CXCL10 and CXCL11 (Li et al., 2020; Wauters et al., 2021) were also highly upregulated at 2DPI (**Figure 3A**). At 7DPI, the robust IFN response in inoculated fLX had subsided (**Figure 3B,D; Figure S5C-E**), but chemokine responses remained evident, highlighting a sustained inflammatory phenotype at the site of infection. Notably, evidence of an IFN systemic response was seen in the contralateral graft at 7DPI (**Figure 3C,D; Figure S5C,D**), despite no evidence of viral replication (**Figure 1P**), providing evidence of transmission of antiviral signals or pathogen-associated molecular motifs (i.e. viral RNA) to contralateral grafts via the peripheral blood. Further, chemokine mRNAs were not upregulated in contralateral grafts (**Figure 3C**), underscoring a link between viral infection, high chemokine expression, persistent inflammation and histopathology. A QIAGEN Ingenuity Pathway Analysis (IPA) supported evidence of prolonged inflammation in inoculated but not in contralateral NRG-L fLX (**Figure 3E**), as many processes involving pro-inflammatory cytokines and mediators (IL-2, IL-7, IL-17, chemokine signaling, T-cell signaling pathways, etc.) were only observed in inoculated fLX. Notably, IPA suggested a key role of the human resident hematopoietic compartment in these inflammatory processes.

**Figure 3.**
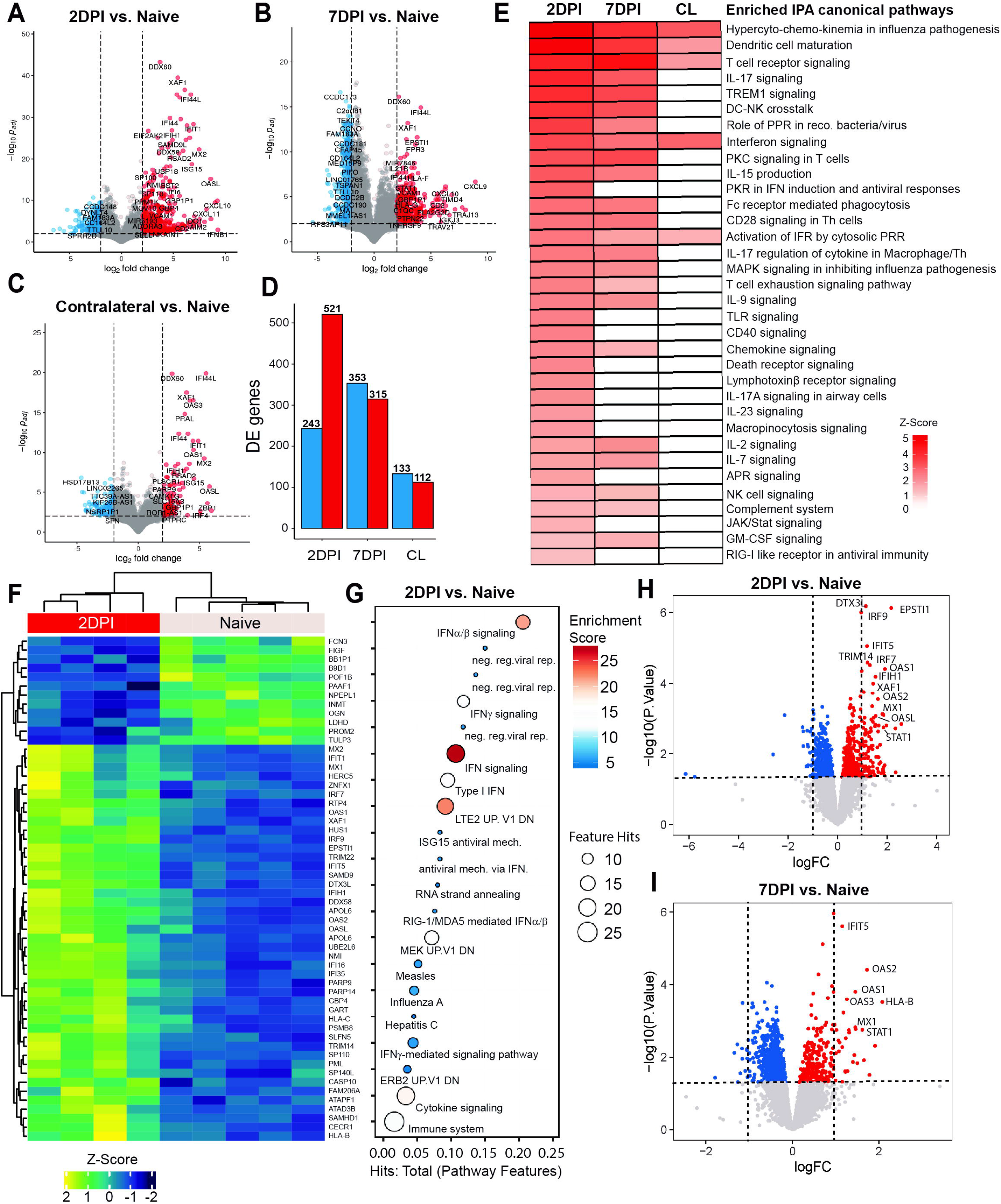
Type I IFN responses and persistent inflammation in inoculated fLX of NRG-L mice. See also Figure S5 and Supplemental Item 3-5. **A-C.** Differentially expressed transcripts in inoculated (A: 2DPI; B: 7DPI) and contralateral non-inoculated fLX (C: 7DPI) following SARS-CoV-2 infection (10^6^ PFU) in comparison to naïve fLX. Transcripts with *p_adj_*≤0.05 and with log_2_ fold change≥2 are considered significantly up- (red) or downregulated (blue). Naïve, n=3; 2DPI, n=4; 7DPI, n=6; CL/Contralateral, n=3. **D.** Number of differentially up-(red) or downregulated (blue) genes per time point (2/7DPI) and infection settings (inoculated/CL). Naïve, n=3; 2DPI, n=4; 7DPI, n=6; CL/Contralateral, n=3. **E.** Enriched IPA canonical pathway analysis (Qiagen) in inoculated fLX (2/7DPI) and in non-inoculated contralateral fLX (CL). Color intensity is proportional to Z-Score. Naïve, n=3; 2DPI, n=4; 7DPI, n=6; CL/Contralateral, n=3. **F.** Cluster heatmap representing protein up- (Z-score>0) and downregulated (Z-Score<0) in NRG-L fLX at 2DPI (n=4; 10^6^ PFU of SARS-CoV-2) in comparison to naïve fLX (n=5). **G.** Protein pathway enrichment analysis at 2DPI (n=4) in comparison to naïve fLX (n=5). Level of enrichment for each protein pathway is colored coded. **H-I.** Volcano plots displaying differentially expressed proteins at 2DPI (H) and 7DPI (I) in inoculated fLX of NRG-L mice. Proteins with *p*≤0.05 (horizontal dashed line) and with logFC≥1 or ≤-1 (vertical dashed lines) are considered significantly up- or downregulated respectively. Naïve, n=5; 2DPI, n=4.

Proteomic analysis were consistent with our transcriptomic findings, and revealed the induction of a canonical, robust type I IFN response at 2DPI which resolved at 7DPI (**Figure 3F-I; Figure S5F-H; Supplemental Item 4**). We could not however detect any upregulation of IFNB1 and IFNL1 or CXCL9-11, as mass spectrometry is known to be a suboptimal method for quantifying cytokine and chemokine expression. Phospho-proteomic signatures also supported the induction of type-I IFN responses (**Figure S6; Supplemental Item 5**), especially at 2DPI and suggested an interesting link between SARS-CoV-2 infection and PML-nuclear bodies (PML-NBs). Phosphorylation of PML-NBs associated proteins SP100 and SP110 was significantly increased at both 2DPI and 7DPI (**Figure S6C,D**), both of which have been previously identified as antiviral restriction factors (Sengupta et al., 2017; Stepp et al., 2013). Two IFN-induced proteins, XAF1 and SAMHD1, showed significant increases in phosphorylation at both 2DPI and 7DPI.

Altogether, persistence of SARS-CoV-2 replication and severe histopathology associates with prolonged inflammation, as well as with resolving an ineffective ISG responses in NRG-L mice.

### fLX in HNFL mice have increased human hematopoietic reconstitution

As we hypothesized that a major cause of the inability of fLX to effectively control infection was the lack of critical hematopoietic effectors able to strengthen and balance antiviral responses, we decided to investigate whether a mature human hematopoietic compartment in fLX would protect fLX from infection. Although human hematopoietic stem cell (HSC) engraftment in NRG mice has been shown to result in robust human lymphoid chimerism, myeloid lineages and NK cell frequencies remain low, which impairs antiviral innate immune responses and T-cell mediated immunity (Douam et al., 2018). Addressing these caveats, we previously reported that NRG mice with a targeted disruption of fetal liver kinase 2 expression (NRG-Flk2^-/-^ or NRGF) display a selective expansion of the human myeloid and NK cell compartment upon adenovirus-mediated delivery of human FMS-related receptor tyrosine kinase 3 ligand (hFlt3LG), the ligand of Flk2 (Douam et al., 2018). We found that these mice were able to mount enhanced innate and T-cell responses against the yellow fever vaccine strain 17D (Douam et al., 2018). Therefore, we aimed here to engraft NRGF mice with pairs of fLX similarly to NRG mice, prior to injection of allogeneic human hematopoietic stem cells 3-5 weeks post fLX engraftment (**Figure 4A**).

**Figure 4.**
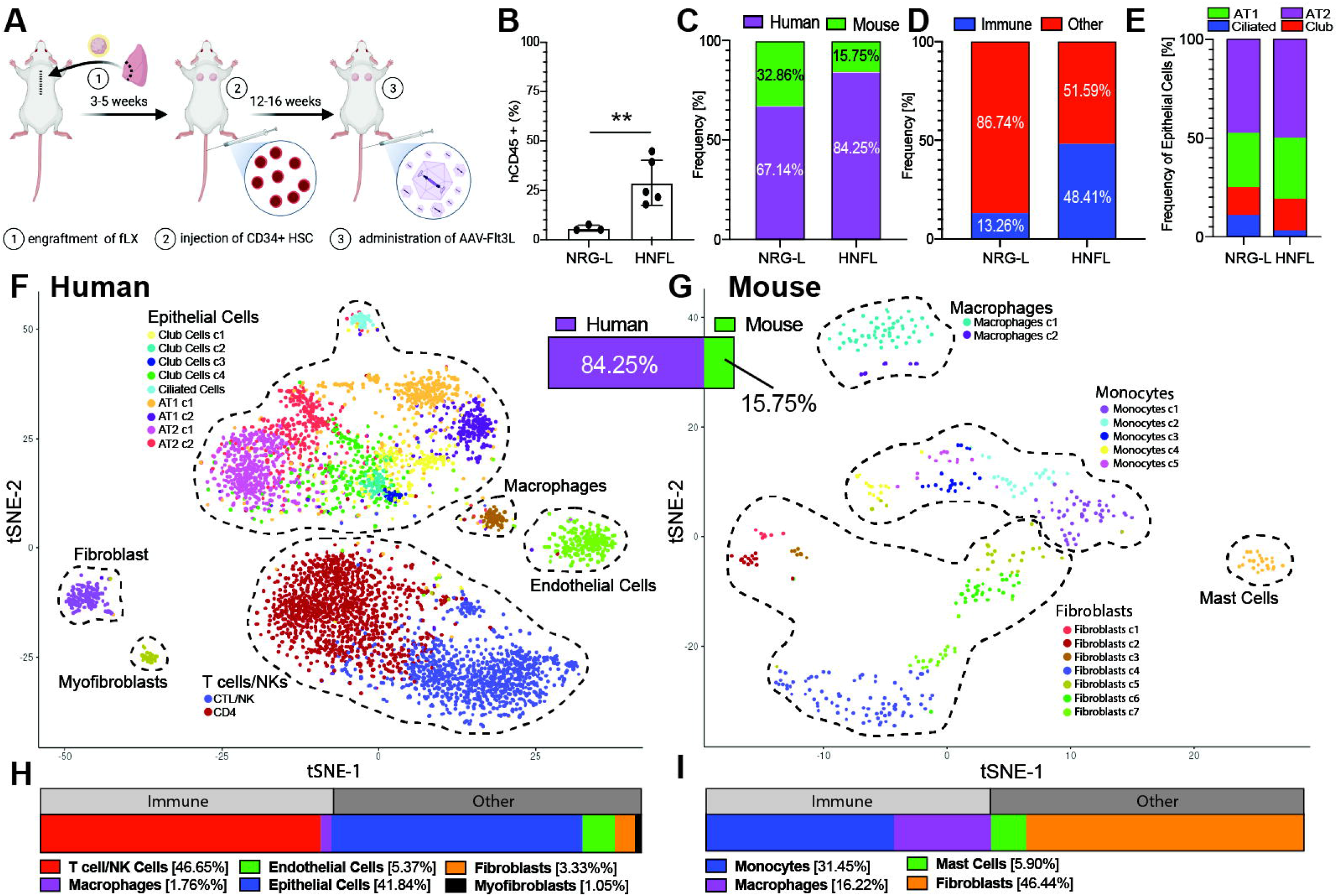
Generation and hematopoietic characterization of fLX in HNFL mice. See also Supplemental Item 1. **A.** Schematic representation of the procedure to generate HNFL mice. Rendered by BioRender. **B.** Frequencies of human CD45+ cells within total CD45+ cells (mouse + human) in naïve fLX of NRG-L and HNFL mice. n=3-5. Mean±SEM, Welch’s t test \*\**p*≤0.01 **C.** Fraction (%) of the human (purple) and mouse (green) cellular compartment in fLX of NRG-L (3 fLX, 15,059 cells) and HNFL (2 fLX, 5,567 cells) as determined following scRNAseq analysis. **D.** Frequencies of the human hematopoietic fraction within the entire human cellular compartment in naïve fLX of NRG-L (3 fLX, 9,968 cells) and HNFL (2 fLX, 5160 cells) as determined by scRNAseq analysis. **E.** Frequencies of AT1 (green), AT2 (purple), ciliated cells (blue) and club cells (red) within the human epithelial compartment in naïve fLX of NRG-L (3 fLX, 2,650 cells) and HNFL (2 fLX, 2,159 cells) as determined by scRNAseq analysis. **F-G.** t-SNE plot of the human (F: 2 fLX, 5,160 cells) and mouse (G: 2 fLX, 407 cells) compartment in naïve HNFL fLX. Relative representation of each compartment within fLX is indicated as the percentage between the two t-SNE plots. Cell subset frequencies within each compartment (H: human; I: mouse) are shown below the respective t-SNE plot.

Twelve to sixteen weeks post HSC engraftment, we injected mice with 2x10^11^ copies of adeno-associated virus (AAV) encoding for hFlt3LG to promote expansion of the myeloid compartment (yielding HIS-NRGF/Flt3LG-L, or HNFL), and analyzed the human hematopoietic reconstitution of fLX 14 days post AAV injection. Using flow cytometry, we found that frequencies of human CD45^+^ were significantly increased in fLX of HNFL in comparison to NRG-L mice (**Figure 4B**). Single-cell RNA sequencing (scRNAseq) confirmed an increased frequency of overall human cells in fLX of HNFL mice (84.25% v. 67.14% in NRG-L) (**Figure 4C**), and this increase was haematopoietically mediated (48.41% human hematopoietic cells v. 51.59% non-hematopoietic; 13.26% v. 86.74% in NRG-L) (**Figure 4D**).The lung epithelial compartment remained intact in fLX of HNFL upon HSC engraftment, with an AT2 sub-compartment encompassing 50.2% of total epithelial subsets vs. 47.7% in fLX from NRG-L mice (**Figure 4E; Supplemental Item 1**). Human T-cell and NK cells represented the largest cellular compartment in fLX of HNFL mice (46.65% of all lineages/subsets) (**Figure 4F,H**). Consistent with the lack of Flk2 expression, the mouse myeloid compartment in fLX of HNFL was reduced (15.75%) over NRG-L mice, and was mainly composed of monocytes, mast cells and macrophages subsets (**Figure 4C,G,I**). We identified alveolar macrophages (1.76%) as the sole human myeloid subset in fLX beyond lymphoid subsets (**Figure 4F; Figure S7A**). We found that the presence of macrophages was a defining characteristic of the fLX HNFL when compared to NRG-L mice, as back-to-back proteomic analysis of naïve fLX between NRG-L and HNFL showed significant upregulation of many macrophage markers such as HCK, SLC9A9, and CD163 (**Figure S7B**). Consistently, quantification of CD68+ cells using 6-color imaging and quantitative image analysis confirmed the significantly higher engraftment of fLX for CD68^+^ macrophages in HNFL mice (**Figure S7C**). The majority of CD68^+^ cells were located within airspaces, but were also observed in the interstitium (**Figure 5A-C**). Altogether, our findings show that the HNFL model represents a promising platform to explore the role of the human myeloid compartment in regulating SARS-CoV-2 infection in the human lung.

**Figure 5.**
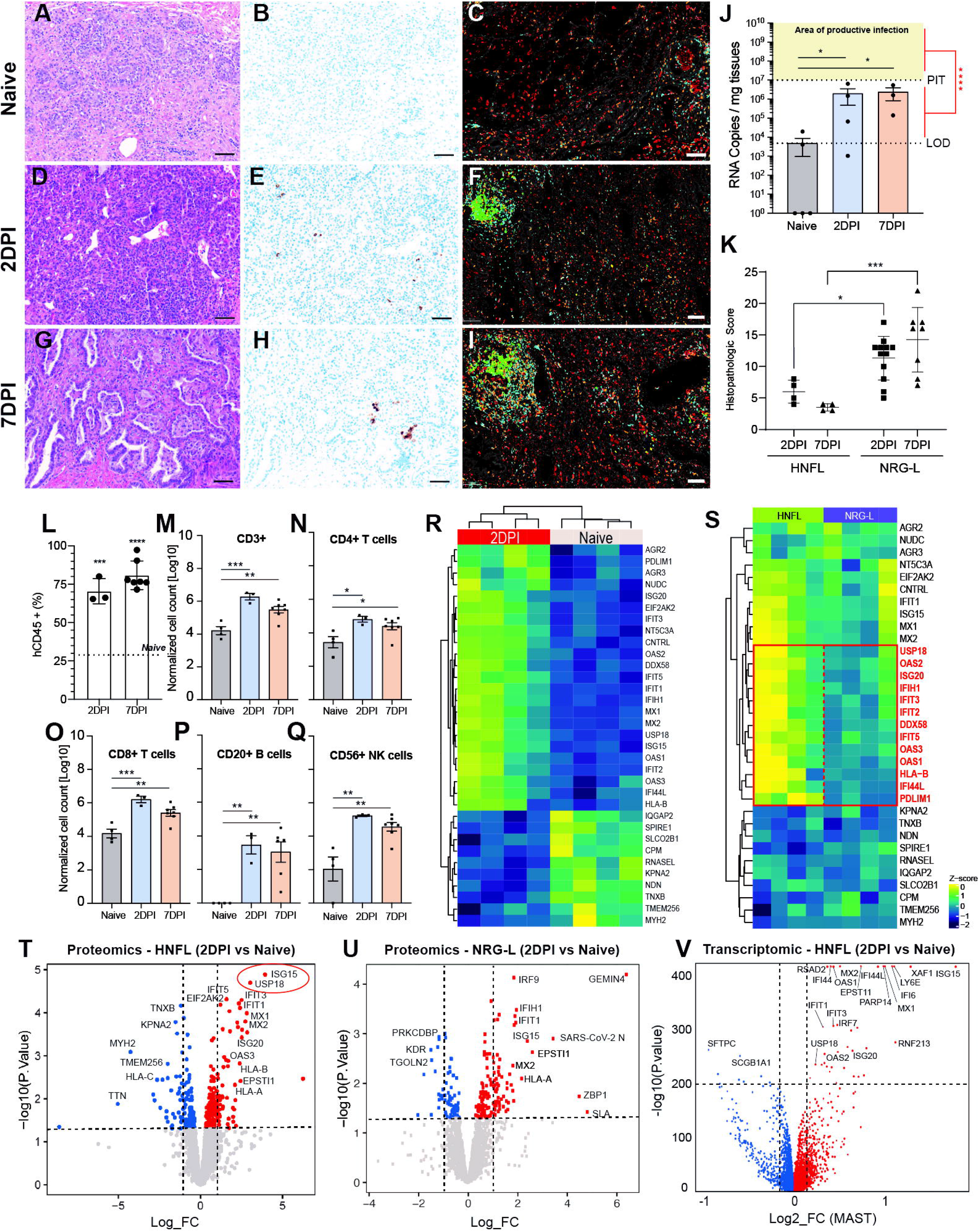
HNFL mice are protected from SARS-CoV-2 infection. See also Figure S5-6 and Supplemental Items 6-7. **A-I.** Representative H&E staining (A,D,G), SARS-CoV-2 Spike IHC (B,E,H) and 6-color IHC (C,F,I: Yellow, SARS-CoV-2 Spike; Cyan, human CD3e; Green, human CD20; Orange, human CD8; Red, human CD68; Grey, DAPI) on naive (A,B,C), or inoculated (10^6^ PFU) fLX tissue sections (B,E,F: 2DPI; G,H,I: 7DPI) from HNFL mice. A,B, D, E, G, H: 200x, scale bar=100µm; C, F, I: 100x, scale bar=200 µm. **J.** SARS-CoV-2 viral RNA quantification by RT-qPCR in inoculated fLX (10^6^ SARS-CoV-2 PFU) of HNFL mice at 2DPI and 7DPI (n=3-5 fLX). Limit of detection (LOD) is shown as a dotted line and represents the mean RNA copies/mg tissues in naïve fLX (n=5). Significance between 2DPI and 7DPI viral load values and productive infection (yellow area) was calculated by running a Kruskal-Wallis test (red line and asterisks) on all pooled 2DPI2 + 7 DPI values from HNFL (n=7) versus all pooled 2DPI+7DPI viral load values from NRG-L mice that were above PIT (n=14). Mean±SEM, Kruskal-Wallis test, \**p*≤0.05. \*\*\*\**p*≤0.001. PIT, productive infection threshold. **K.** Cumulative histopathologic score of inoculated fLX (10^6^ PFU of SARS-CoV-2) from NRG-L and HNFL mice at 2DPI and 7DPI. n=4-12. Mean±SEM, two-way ANOVA \**p*≤0.05 \*\**p*≤0.01. **L.** Frequencies of human CD45+ cells within total CD45+ cells (mouse+human) in fLX of HNFL mice at 2DPI and 7DPI. A dotted line represents the mean frequency of CD45+ cells in naïve fLX from HNFL mice. n=3-7. Mean±SEM, one-way ANOVA \*\**p*≤0.01, \*\*\**p*≤0.001 over naïve fLX (Figure 4B). **M-Q.** Normalized cell count (number of cells analyzed for a given subset*[total fLX cell count/total cells analyzed]) of CD3+ T cells (M), CD3+ CD4+ T cells (N), CD3+ CD8+ T cells (O), CD20+ cells (P) and CD3- CD20- CD33- CD56+ (Q) in naïve fLX or infected (2DPI and 7 DPI). n=3-7. Mean±SEM, one-way ANOVA. \**p*≤0.05, \*\**p*≤0.01, \*\*\**p*≤0.001. **R.** Cluster heatmap representing proteins significantly (*p*≤0.05) up- (z-score>0) and downregulated (z-Score<0) in HNFL fLX at 2DPI (10^6^ PFU of SARS-CoV-2) in comparison to naïve HNFL fLX. Naïve, n=4; 2DPI, n=4. **S.** Semi-cluster heatmap representing relative differential expression of a set of selected proteins in HNFL fLX and NRG-L at 2DPI (10^6^ PFU of SARS-CoV-2) following side-by-side analysis by mass spectrometry run. Protein significantly (*p*≤0.05) up- (z-score>0) and downregulated (z- Score<0) are labelled in red. HNFL, n=4; NRG-L, n=4. **T-U.** Volcano plots displaying differentially expressed proteins in HNFL (T) or NRG-L mice (U) fLX at 2DPI following side-by-side analysis by single mass spectrometry run. Proteins with *p*≤0.05 (horizontal dashed line) and with logFC≥1 or ≤-1 (vertical dashed lines) are considered significantly up- or downregulated respectively. Naïve, n=4; 2D PI, n=4. **V.** Significantly (*p*≤0.05) differentially expressed transcripts (upregulated, red; downregulated, blue) in inoculated HNFL fLX at 2DPI following SARS-CoV-2 infection (10^6^ PFU) in comparison to naïve HNFL fLX. Fold changes were derived from scRNAseq datasets and were computed using MAST. Transcripts with *p*≤10^-200^ (horizontal dotted line) and with log_2_ fold change≥0.2 or ≤- 0.2 (vertical dotted lines) are highlighted. Naïve, n=2; 2DPI, n=3.

### fLX in HNFL mice are protected from SARS-CoV-2 infection

In striking contrast to NRG-L mice, inoculation of HNFL fLX with SARS-CoV-2 (10^6^ PFU) one-week post AAV-Flt3LG injection resulted in the absence of productive infection, as assessed by low to no SARS-CoV-2 Spike immunoreactivity (**Figure 5A-I**) and by the detection of viral RNA copies per mg of tissues significantly below PIT (**Figure 5J**). Importantly, the lack of Flk2 expression did not affect fLX susceptibility to infection (**Figure S7D**), underscoring that the protective phenotype was not mediated by a reduced mouse myeloid compartment. Despite very limited infection events, spike antigen distribution in HNFL mirrored that observed with NRG-L mice, confined to the cytoplasm of viable-looking AT2 pneumocytes and bronchiole epithelium (**Figure 5E,H**), but with little to no discernible cytopathic effect or neighboring intra-airspace necrosis.

Consistently, cumulative histology scores were significantly decreased in HNFL mice at 2DPI (*p*=0.01) and 7DPI (*p*=0.0002) when compared to NRG-L (**Figure 5K; Figure S7E**). HNFL fLX mice showed decreased syncytial cells and intra-airway necrosis at 2DPI and 7DPI, respectively when compared to NRG-L fLX. Hemorrhage and influx of neutrophils were also significantly decreased at both 2DPI and 7DPI for HNFL fLX compared to NRG-L fLX.

Taken together, these findings suggest that human hematopoietic engraftment in HNFL may promote an effective host response in fLX that prevents the establishment of persistent SARS-CoV-2 infection, and protects tissues from diffuse alveolar damage.

### HNFL mice mount a potent and balanced type I IFN response against SARS-CoV-2

Human CD45^+^ frequency (**Figure 5L**) and human T cell, B cell, and CD56^+^ normalized cell counts significantly increased in fLX of HNFL mice following virus infection (**Figure 5M-Q; Figure S7F**), supporting evidence of successful viral inoculation despite effective control of viral replication. Consistently, a striking feature of comparative 6-color immunostaining before and after infection was the formation of CD20^+^ B cell lymphoid aggregates post infection (**Figure 5F,I**).

To provide further evidence of active host responses, we conducted proteomic analysis of HNFL fLX cell lysates prior to and after infection. Proteomic analysis revealed that in HNFL fLX, infection resulted in the induction of a significant type I IFN response at 2DPI, notably characterized by the upregulation of ISG15, USP18, IFIT1, IFIT3, and MX1-2 (**Figure 5R; Supplemental Item 6**). Notably, side by side proteomic analysis at 2DPI indicated that HNFL fLX displayed a stronger upregulation of a broad set of ISGs in comparison to NRG-L fLX (**Figure 5S**), including USP18, OAS1-3, ISG20, IFIH1, IFIT2-3, DDX58, IFIT5, or IFI44L. Consistently, SARS-CoV-2 N was upregulated in NRG-L fLX but not in HNFL fLX. A striking and specific feature of the proteomic response in HNFL fLX was that it was dominated by the upregulation of the USP18-ISG15 axis, unlike NRG-L fLX, which did not display any significant USP18 upregulation (**Figure 5T-U**). USP18 is the main ISG15 isopeptidase and a well-known inhibitor of type I IFN signaling (Honke et al., 2016). ISG15 is known to promote USP18-mediated inhibition of type I IFN signaling by stabilizing USP18 activity and preventing its proteasomal degradation (Zhang et al., 2015), underscoring the role of ISG15 as a negative regulator of type I IFN responses when co-upregulated with USP18. Consistent with the function of the USP18-ISG15 axis, phospho-proteomic analysis confirmed that STAT1 was significantly phosphorylated in NRG-L, but not in HNFL fLX (**Figure S7G,H; Supplemental Item 7**). Bulk transcriptomic analysis of HNFL fLX at 2DPI confirmed significant upregulation of the USP18-ISG15 axis (FDR=6.69e-235 and 0, respectively), as well as the strong upregulation of many other ISGs such as RSAD2, IFI44/L, OAS1, MX1-2, IFI6, LYE6 (FDR=0) (**Figure 5V**). Unlike NRG-L fLX, no chemokines were among the top upregulated hits (Log2FC≥0.2 and FDR≤1e-200). IFNB1/L1 was not detectable likely due to the already extensive viral clearance observed at 2DPI.

Of note, SFTPC was the most downregulated transcript at 2DPI (-0.9 Log2FC; FDR=4.42e-232). This finding is consistent with previous studies reporting a loss of AT2 program/compartment upon SARS-CoV-2 infection (Delorey et al., 2021), and further supports the relevance of the HNFL mouse model to study immune correlates of protection against SARS-CoV-2. At 7DPI, ISG and SFTPC (-0.5 Log2FC; FDR=1e-109) were found to be returning to naïve fLX expression levels (**Figure S7I**), which is consistent with the rapid resolution of infection of fLX. Altogether, the superior ISG upregulation and absence of a strong proinflammatory response observed in HNFL fLX in comparison to NRG-L fLX underscores a robust association between effective control of viral infection and limited histopathology, and the development of a potent and balanced antiviral response against SARS-CoV-2 infection.

### Macrophages activate and differentiate upon SARS-CoV-2 infection in HNFL fLX

Finally, we used scRNAseq to define which cellular compartment(s) mediate this potent ISG response associated with tissue protection. scRNAseq analysis confirmed significant hematopoietic infiltration in HNFL fLX at 2DPI (78.89% human cell vs. 48.41% in naïve fLX) and progressive resorption of such infiltration by 7DPI (68.52% human cells) (**Figure 6A-E; Supplemental Item 1**). The major subsets mediating hematopoietic expansion in infected fLX were macrophages (Naïve, 1.76%; 2DPI, 18.39%; 7DPI, 20.49%) and B-cells (Naïve, 0%; 2DPI 7.24%; 7DPI, 3.31%) (**Figure 6F**). Consistent with our findings above, the size of the AT2 sub-compartment within the epithelial compartment was reduced at 2DPI (12.7%) but restored at 7DPI (45.5% vs. Naïve, 50.2%) (**Figure 6G)**, supporting evidence for the successful resolution of infection and progressive restoration of tissue homeostasis. Given the significant increase in macrophages in infected fLX and their known functions in controlling viral infection, we decided to focus our dataset analysis on this cellular compartment. At 2DPI, we identified two differentially polarized CD14^+^ CD68^+^ activated macrophage subsets; one had a pro-inflammatory phenotype (herein referred to as activated inflammatory macrophages, [AIM]) defined as CD14^+^ CD68^+^ IL1β^+^ and which displayed moderate ISG expression and low IL10 expression. The other had a regulatory phenotype (subsequently referred to as activated regulatory macrophages, [ARM]) defined as CD14^+^ CD68^+^ IL1β^+^ C1QA^+^ and which displayed high ISG and moderate IL10 expression (**Figure 6H-I**). Similar macrophage subsets were observed at 7DPI (**Figure S7J**). Activated macrophage clusters were significantly associated with moderate expression of several chemokines at 2DPI, including CCL2-4, CCL8 and CXCL10 which was consistent with monocyte recruitment and macrophage infiltration into fLX upon infection. Expression of CCL13 and CCL18, considered to be pro- and anti-inflammatory chemokines, respectively, (Mendez-Enriquez and Garcia-Zepeda, 2013; Schraufstatter et al., 2012), were significantly associated with the ARM cluster only, underscoring the role of ARM as a major regulatory subset.

**Figure 6.**
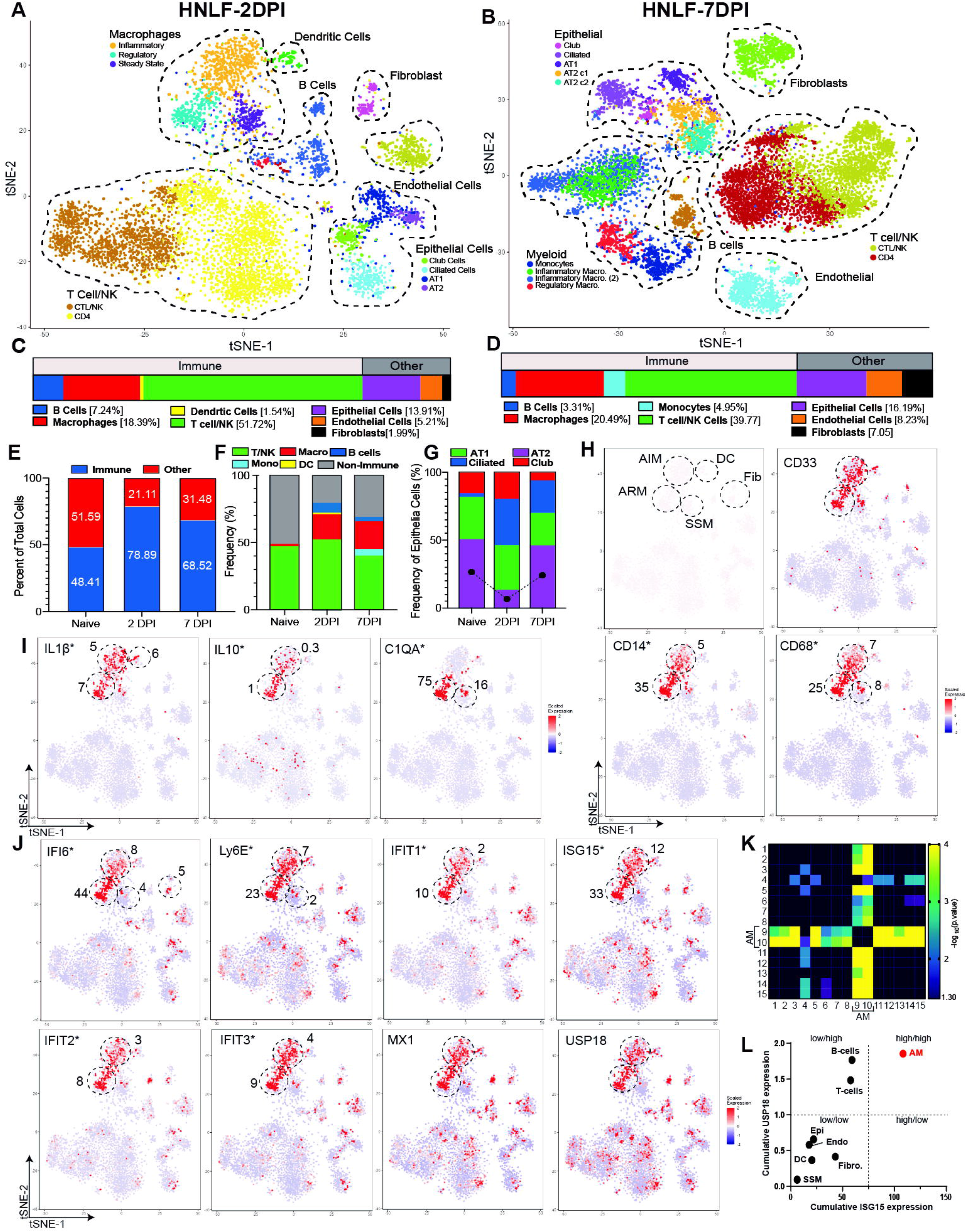
Macrophages are the major mediator of the ISG response observed in HNFL mice. See also Figure S7. **A-D.** t-SNE plot of the human compartment in fLX of HNFL mice at 2DPI (A: 3 fLX, 6,736 cells) and 7DPI (B: 3fLX, 11,269 cells). Detailed cell subset frequencies for each time point (C: 2DPI; D: 7DPI) are shown below the respective t-SNE plot. **E.** Frequencies of the human hematopoietic fraction (immune) within the entire human cellular compartment in naïve HNFL fLX (2 fLX, 5160 cells) or after inoculation (2DPI: 3 fLX, 6,736 cells; 7DPI: 3fLX, 11,269 cells). **F.** Frequencies of different human hematopoietic lineages within the entire human cellular compartment (2 fLX, 5,160 cells) in naïve HNFL fLX or after inoculation (2DPI: 3 fLX, 6,736 cells; 7DPI: 3 fLX, 11,269 cells). **G.** Frequencies of AT1 (green), AT2 (purple), ciliated cells (blue) and club cells (red) within the human epithelial compartment in naïve HNFL fLX (2 fLX, 2,159 cells) or after inoculation (2DPI: 3 fLX, 937 cells; 7DPI: 3fLX, 1,825 cells). A dotted line symbolizes the variation of the size of the AT2 compartment upon infection. **H-J.** t-SNE plots displaying clustered and scaled expression of several transcripts coding for human cell surface markers and ISGs in the human compartment of HNFL fLX at 2DPI. (H) Myeloid markers CD33, CD14 and CD68. (I) Inflammatory and regulatory markers IL1β, IL10 and C1QA. (J) ISG IFI6, Ly6E, IFIT1-3, MX1, ISG15 and USP18. Clusters of interest are indicated with a dotted circle, and each cluster name is shown in the top left plot of panel H (AIM, activated inflammatory macrophages; ARM, activated regulatory macrophages; SSM, steady-state macrophages; DC, dendritic cells; Fib, fibroblasts). Cluster defining genes (i.e., whose expression level is significantly associated with a given cluster) have their name followed by an asterisk, and Log2FC values are indicated near the corresponding cluster(s). n=3 fLX, 6,736 cells. **K.** Statistical association between ISG upregulation and cluster identity. Mean scaled expression of a pool of 15 ISGs (DDX58, IFI44L, ISG15, OAS1-3, IFIT1-3, IFIT5, USP18, LyE6, IFI6, MX1-2) was compared across all the human scRNAseq clusters at 2DPI (n=15). Statistically significant differences in ISG expression between clusters was calculated using a two-way ANOVA test, and p values (-Log_10_(p.value)) are reported as heatmap. *p*≥0.05 (-Log10[0.05]=1.30) are shown in black and are considered non-significant. Activated macrophage (AM) clusters are indicated (clusters 9, inflammatory macrophages; cluster 10, regulatory macrophages) **L.** Differential co-expression of the USP18-ISG15 axis across human lineages in infected fLX. Cumulative scaled expression of USP18 and ISG15 was calculated for all human lineages in fLX at 2 DPI (Epithelial cells, Epi: 4 clusters; B-cells: 3 clusters; T-cells: 2 clusters; Endothelial cells, Endo: 1 cluster; Fibroblasts, Fibro: 1 cluster; Dendritic cells, DC: 1 cluster; Steady-state macrophages, SSM: 1 cluster; Activated macrophages, AM: 2 clusters) and plotted on a X/Y axis with X and Y corresponding to ISG15 and USP18 expression respectively. Four categories of co-expression were identified (low/low, low/high, high/low, and high/high) and are delimited by dashed lines.

### Activated macrophages mediate potent IFN-mediated antiviral responses against SARS-CoV-2

When analyzing the cellular distribution of top upregulated ISGs from our proteomic and transcriptomic HNFL datasets at 2DPI, we found that activated macrophages ubiquitously expressed every single ISG examined (n=15) (**Figure 6J; Figure S7K)**, and were the dominant source of expression of these ISGs across the entire fLX human compartment (**Figure 6K**). Some of these ISGs, including ISG15, IFI6, LY6E and IFIT1-3, displayed levels of expression that statistically associated with activated macrophage clusters (**Figure 6J, K**). Importantly, activated macrophages also represented the dominant source of USP18-ISG15 co-expression (**Figure 6L**) strengthening their key role in regulating antiviral responses during infection. Strikingly, activated macrophage clusters across both time points (n=5) were found to be the major carriers of viral RNA (*p*=0.001) versus all other clusters (n=24), suggesting a potential association between the dominant macrophage-mediated ISG response (**Figure S7L**) and direct activation by viral RNA.

Altogether, our findings highlight infiltrating macrophages as the central mediator of the potent and well-balanced protective IFN response at play in HNFL fLX, and suggest that direct activation by viral RNA might be a key element triggering that process.

## DISCUSSION

For the majority of healthy individuals below middle age, COVID-19 remains a mild to asymptomatic disease (Nikolai et al., 2020; Wu and McGoogan, 2020). Despite this, most studies have focused on immunological processes driving severe COVID-19 cases, using data from human patients (Delorey et al., 2021; Rendeiro et al., 2021). To enhance our understanding of the cellular and molecular mechanisms driving protective immune responses in the respiratory tract of asymptomatic and mild COVID-19 patients, investigations need to go beyond human patient studies due to their inherent limitations.

Immunodeficient mice engrafted with fLX have been previously reported to be highly permissive to coronavirus infection (Wahl et al., 2021). Such *in vivo* platforms carry substantial advantages over other *in vivo* models (including humans) in a way that enables investigations into coronavirus infection within a human lung environment over time and in controlled experimental settings. However, given our common understanding that the quality of immune responses during SARS-CoV-2 infection defines disease outcome, the lack of significant human hematopoietic reconstitution in these models has precluded their use for investigating human immune responses against coronavirus infection. Here, we report that NRG-L mice singly engrafted with fLX are highly susceptible to SARS-CoV-2 infection. Inoculated fLX were prone to extensive inflammation, which associated with severe histopathological manifestations of disease. In sharp contrast, co-engraftment of fLX and human HSC in HNFL mice resulted in protection against SARS-CoV-2 infection, as well as limited inflammation and histopathology.

Protection associated with the induction of a superior ISG response in HNFL mice as compared to NRG-L mice, which was dominated by the upregulation of the USP18-ISG15 axis – a unique feature of inoculated HNFL fLX. USP18-ISG15 upregulation correlated with the absence of prolonged inflammation and severe histopathology, consistent with its role as a negative regulator of antiviral responses. Inoculation of HNFL fLX with SARS-CoV-2 induced a significant infiltration of macrophages in fLX. We identified infiltrating activated macrophages as the dominant mediator of the prominent type I IFN response observed in infected fLX, as well as the major compartment co-expressing USP18 and ISG15. Finally, we also identified activated macrophages to be the primary carriers of viral RNA. Altogether, our work highlights lung infiltrating macrophages as a multi-faceted hematopoietic subset mediating front-line antiviral responses against SARS-CoV-2 while also safeguarding lung tissue integrity. Our findings provide unique molecular correlates of lung tissue protection during SARS-CoV-2 infection, and propose a unique model in which macrophage-mediated ISG responses would promote a systemic antiviral response against SARS-CoV-2, and tight control of these responses via the USP18-ISG15 axis would prevent excessive inflammation and severe histopathology.

Our study suggests a working model (**Figure 7**) in which lung resident macrophages (e.g. alveolar macrophages) would become locally activated upon detection of SARS-CoV-2 RNA, and activation would promote monocyte infiltration via regulated secretion of specific inflammatory chemokines (CCL2/4/8, CXCL10). Monocyte infiltration would be followed by macrophage differentiation, activation, and polarization toward inflammatory and regulatory phenotypes. These two subsets of macrophages would then promote a systemic antiviral state across the epithelial compartment in a coordinated fashion, enabling rapid suppression of viral spread and replication. The USP18-ISG15 axis would tightly control those responses in order to protect the tissue from excessive inflammation. In contrast, in NRG-L mice, the pro-inflammatory response mediated by resident human macrophages following detection of viral RNA would remain a “call in the dark.” Indeed, the absence of human hematopoietic engraftment in NRG-L mice preclude the possibility for any human monocyte recruitment into the fLX, therefore preventing the induction of a stronger, systemic antiviral response that could rapidly and effectively clear infection. Additionally, the absence of infiltrating macrophages would allow the initial inflammatory response to go unhindered between the resident human lymphoid and myeloid compartment of the fLX, promoting diffuse alveolar damage and other histopathological manifestations of disease while further dampening effective antiviral responses. This model is supported by our observation of the antiviral responses in contralateral fLX. While contralateral fLX mounted a robust type I IFN response, they did not display any histopathological manifestations of disease and prolonged inflammation. This finding is consistent with the idea that fLX resident cells need to come in contact with a sufficient amount of infectious viral particles or viral RNA to trigger a damaging inflammatory loop. In the case of the contralateral grafts, the amount of viral RNA transferring from the inoculated graft was likely too low.

**Figure 7.**
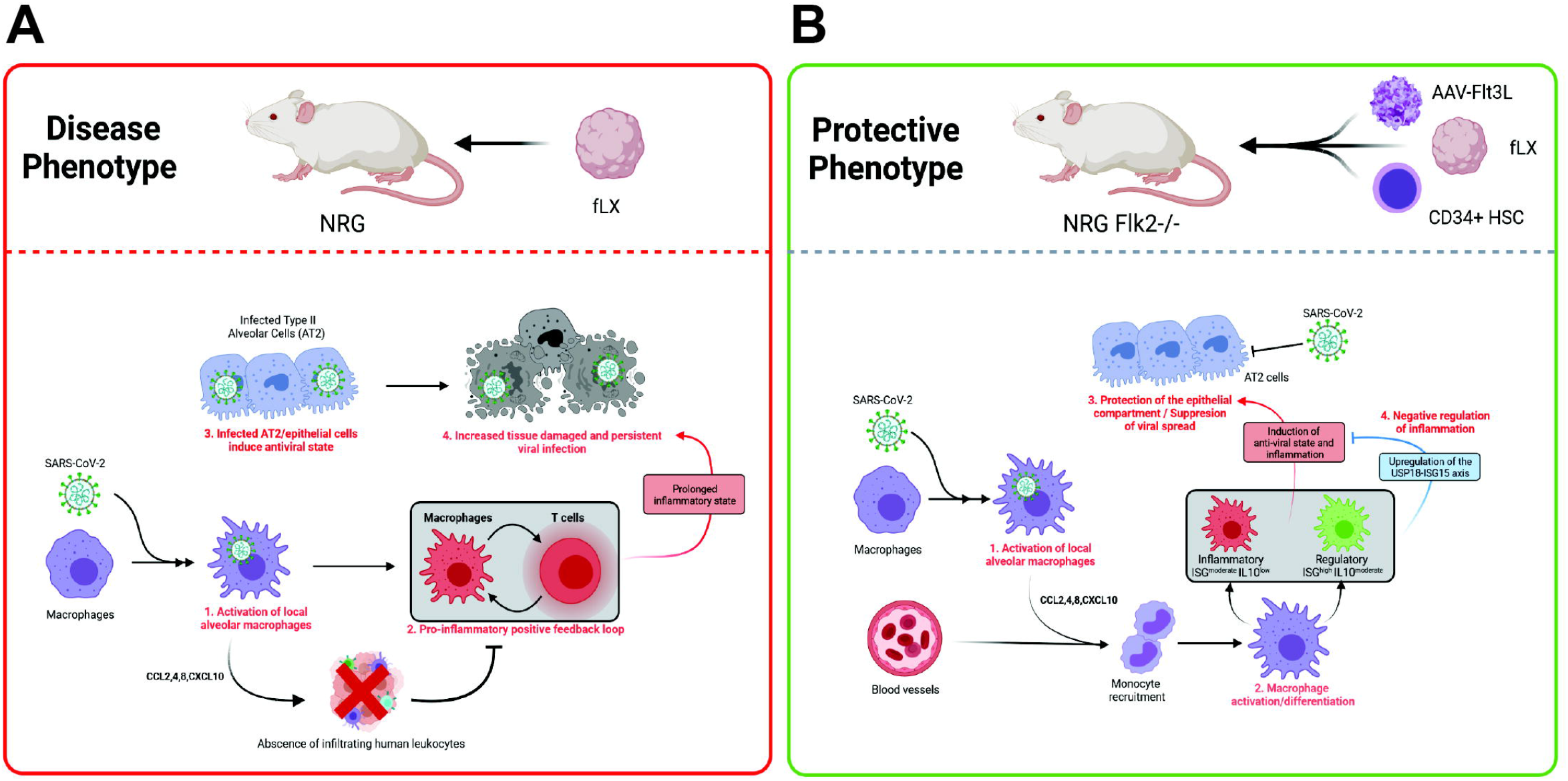
Working model: Macrophages as central regulator of lung tissue protection upon SARS-CoV-2 infection. **A.** Disease model: In NRG-L, macrophage infection results in pro-inflammatory chemokine secretion (1) and in a pro-inflammatory feedback loop (2) that goes unhindered by the absence of human leukocyte infiltration in infected fLX. The absence of macrophage infiltration does not allow for the establishment of a type-I interferon response able to effectively support the antiviral response mediated by the type II pneumocyte (AT2) compartment (3), therefore preventing effective viral clearance. Persistent viral infection and prolonged inflammation over time induce diffuse alveolar damage (4) and other histopathological features observed in the lung of severe cases of COVID-19. **B.** Protective phenotype: In HNFL mice, macrophages are activated upon SARS-CoV-2 infection (1) and promote the recruitment of monocytes via chemokine production including CCL2, CCL4, CCL8 and CXCL10. Monocyte differentiation into macrophages (2), and further polarization into inflammatory and regulatory phenotypes, promotes a systemic antiviral state within the fLX (3). Upregulation of the USP18-ISG15 axis allows for a tightly balanced inflammatory response that preserves lung tissue integrity during the antiviral response (4). Imaged by BioRender.

The resemblance between the histopathological manifestations observed in NRG-L fLX and those observed in severe COVID-19 patients suggest that NRG-L may indirectly recapitulate important disease mechanisms occurring in such patients. Macrophage excessive pro-inflammatory responses have been hypothesized to be an important driver of COVID-19 disease progression (Grant et al., 2021; Merad and Martin, 2020; Rendeiro et al., 2021; Wauters et al., 2021). Therefore, it is conceivable that the human macrophage-mediated prolonged inflammation, and/or the absence of effective regulation of inflammation by infiltrating macrophages in NRG-L, act as a surrogate of the immunological dysregulations observed in severe COVID-19 patients. As our understanding of how macrophages contribute to COVID-19 disease in humans so far is largely speculative, further work is now needed to experimentally challenge and validate our working model, which will be the focus of our follow-up studies. Our HNFL model opens up unprecedented avenues to do so, and to evaluate specifically the impact of macrophages on viral replication, inflammation, and histopathology in a human tissue context and in controlled experimental settings. Such future investigations could lay the groundwork for the development of innovative macrophage-targeting immunotherapies against viral respiratory diseases.

Consistently with our working model, several studies have reported the susceptibility of monocyte-derived macrophages or alveolar macrophages to SARS-CoV-2 uptake, but not to active viral replication (Boumaza et al., 2020; Dalskov et al., 2020; Grant et al., 2021; Yang et al., 2020a). Specifically, macrophage exposure to SARS-CoV-2 can result in a strong pro-inflammatory response but attenuated type I IFN response, suggesting that non-infected infiltrating macrophages might be more effective in mediating and regulating systemic antiviral responses than initially infected macrophages (Yang et al., 2020a). However, as macrophage differentiation and antiviral responses depend on their tissue and extracellular environment, it remains challenging to extrapolate such *in vitro* data into an *in vivo* tissue context. Further work is needed to delineate the connection between macrophage infection and macrophage-mediated IFN response and inflammation in HNFL mice.

A unique finding of our study is the strong association between the upregulation of the USP18-ISG15 axis by macrophages and limited histopathology and inflammation. The role of the axis in regulating viral infection in a human tissue context remains elusive. ISG15 is mainly recognized as having antiviral functions, and many viruses have developed ways to escape such functions (Perng and Lenschow, 2018). USP18 is the main ISG15 isopeptidase and can inhibit ISG15-mediated ISGylation and can repress the establishment of an antiviral state (Honke et al., 2016; Ketscher et al., 2015). However, USP18 can also directly inhibit type I IFN signaling by binding to STAT2 (Arimoto et al., 2017) and IFNAR2 (Malakhova et al., 2006). Most importantly, ISG15 is a crucial partner to USP18 in this process (Zhang et al., 2015) by stabilizing it and preventing its proteasomal degradation, highlighting ISG15 as a negative regulator of the type I IFN response. As macrophage-mediated inflammation has been suggested by many human studies to contribute to COVID-19 (Grant et al., 2021; Merad and Martin, 2020; Rendeiro et al., 2021; Wauters et al., 2021), and the USP18-ISG15 axis is not upregulated in NRG-L mice, our data imply that upregulation of the USP18-ISG15 axis acts as an important regulator of macrophage inflammation during infection, keeping positive inflammatory feedback loops in check (e.g., via the downregulation of RIG-I-mediated signaling, for instance) in order to prevent excessive tissue damage during host-mediated antiviral responses. This working model is consistent with previous findings in hamsters, in which STAT2 was found to prevent viral dissemination (Boudewijns et al., 2020), while contributing to excessive tissue inflammation. Altogether, our work sheds unique light on the complex and multifaceted role of USP18 and ISG15 during viral infection *in vivo* and within a human tissue context. Further work is now required to evaluate the impact of these proteins and their proteolytic/non-proteolytic functions in the protective phenotype we observed. Given the human specific nature of the interaction between USP18 and ISG15 (Speer et al., 2016), the HNFL model uniquely positions itself for such investigation over other animal models of SARS-CoV-2 infection.

Our study demonstrates how fLX-engrafted mice can overcome certain limitations associated with human studies and effectively transforms our understanding of human immunity. However, it is worth noting that such models also carry inherent limitations such as imperfect human immune reconstitution, B-cell responses, and/or imperfect cellular architecture of lymphoid tissues. Significant improvements have been achieved over the past few years to develop advanced models displaying enhanced myeloid compartment and innate immune responses (O’Connell and Douam, 2020), some of which were applied in this study. Continuous refinements of these models, such as by implementing genetic strategies to further enhance human immune reconstitution, myeloid and lymphoid functions, and humoral responses, will aid in further increasing our understanding of human immunity to coronavirus infection.

Graft versus host disease (GvHD) also remains an important limitation in accurate modeling of human immune responses in humanized mice (Rongvaux et al., 2013). We previously reported that the NRGF model was particularly resistant to GvHD in comparison to conventional NRG mice due to the depletion of the mouse myeloid compartment in this model (Douam et al., 2018). Despite the fact that HNFL mice were co-engrafted with allogeneic HSC in this study due to limited tissue accessibility, we did not observe any significant signs of human-to-human GvHD in fLX either histologically or by omics analyses. No major evidence of tissue damage in naïve HNFL fLX was observed and no significant epithelial or hematopoietic inflammation was observed by proteomics or scRNAseq analysis. At 7DPI, the restoration of the AT2 compartment and the resolution of inflammatory responses toward baseline level strongly support evidence that the immune signatures observed in fLX are mediated by SARS-CoV-2 infection. In future studies, co-engrafting HNFL with autologous HSCs will help increase the capability of this mouse model. Altogether, our work emphasizes the potential of fetal tissue-engrafted mice to transform our understanding of human immunity to pave the way toward more effective treatments against infectious diseases.

## ACKNOWLEDGMENTS

This work was supported in part by a start-up fund and Peter Paul Career Development Professorship from Boston University (to F.D.), grants from the National Institutes of Health (R01 AI138797, R01 AI107301, R01 AI146917, R01 AI153236 to A.P.; R21 ES032882, K22 AI144050 to F.D.; R01 HL141513, R01 HL139641, R01 AI153613, UL1 TR001430 to M.B.; R01 502088946, R21 AI135517 to J.H.C.), Clinical and Translational Science Awards (grant UL1 TR001430) from the National Center for Advancing Translational Sciences of the National Institutes of Health (to F.D., N.A.C. and A.E.), a grant from the National Library of Medicine (R01 LM013154-01 to J.D.C.), the Deutsche Forschungsgemeinschaft (BO3482/3-3, BO3482/4-1 to M.B.), a Research Scholar Award from the American Cancer Society (RSG-15-048-01-MPC to A.P.), a Burroughs Wellcome Fund Award for Investigators in Pathogenesis (101539 to A.P.), Princeton COVID-19 research funds through the Office of the Dean for Research, and Boston University startup funds and an Evergrande MassCPR award (to M.S.). T.T. is a recipient of postdoctoral fellowship awards from the Uehara Memorial Foundation and the JSPS Research Fellowships for young scientists. We thank the Evans Center for Interdisciplinary Biomedical Research at Boston University School of Medicine for their support of the Affinity Research Collaborative on ‘Respiratory Viruses: A Focus on COVID-19’. This work utilized a Ventana Discovery Ultra autostainer that was purchased with funding from a National Institutes of Health SIG grant (S10 OD026983). We thank the Boston University Animal Science Center, and more particularly Corey Nunes and all the NEIDL animal core staff for their outstanding support. We thank Robert LeDesma and Emily Mesev for their critical review of the manuscript. We also thank all the Douam, Ploss, Crossland, Connor, and Emili Lab members, NEIDL members, and members of the department of microbiology, pathology, biochemistry, and medicine at Boston University for their constant support and advice.

## AUTHOR CONTRIBUTIONS

D.K. and F.D. conceptualized the study. D.K., A.K.O., J.T., P.M., R.M.H., T.T., J.H.C., A.E., N.A.C., A.P. and F.D. designed the experiments. D.K., A.K.O., J.T., P.M., R.M.H., T.T., A.R.B., T.R.C., B.B., S.I.G., B.L.H., H.P.G., A.T., A.J.T., E.C., A.S., S.K., K.G., M.S., A.B.B., N.A.C., A.P. and F.D. performed experiments. D.K., A.K.O., J.T., P.M., R.M.H., T.T., S.A.A., B.B., E.B., M.E., M.S., K.P.F., A.K., N.P., J.D.C., J.H.C., A.E., N.A.C., A.P. and F.D. analyzed the data. J.T., R.M.H., S.A.A. and J.D.C. carried out computational analysis. K.P.F., A.K. and N.P. carried out bioluminescence imaging analysis. M.E. carried out electron microscopy analysis. B.R.H., M.B., K.P.F., A.K., and N.P provided access to key resources. E.B., M.B., M.E., M.S., K.P.F., A.K. and N.P. provided conceptual and technical inputs and/or helped with data interpretation. D.K., N.A.C., A.P. and F.D. wrote the manuscript with contributions from all authors.

## METHODS

### Cells and antibodies

VeroE6 cells and AAV-293 cells (ATCC) were grown in Dulbecco’s modified Eagle’s medium (DMEM) supplemented with 10% heat inactivated fetal bovine serum (Bio-Techne, R&D systems, Minneapolis, MN, USA) and 1% (v/v) Penicillin Streptomycin (Thermo Scientific, Waltham, MA, USA).

The following anti-mouse antibodies were used for flow cytometry: from BioLegend (San Diego, CA, USA): CD45-PE-Cy7 clone 30-F11, CD45-PE-Dazzle5 clone 30-F11. The following anti-human antibodies were used for flow cytometry: from BD Biosciences (San Jose, CA, USA): CD45-V500 clone HI30, CD34-FITC clone 581, CD8-FITC clone G42-8, CD11c-allophycocyanin clone B-ly6; from BioLegend: CD56-allophycocyanin-Cy7 clone HCD56, CD33-PerCP-Cy5.5 clone WM53, CD11c-Alexa Fluor 700 clone 3.9, CD123-eFluor450 clone 6H6, CD14-PE-eFluor610 clone 61D3, CD3-FITC clone SK7, CD19-allophycocyanin clone HIB19, HLA-DR-BV510 clone L243, CD3-BV605 clone UCHT1, CD20-BV650 clone 2H7, CD16-BV117 clone B73.1, CD45-BV785 clone H130, CD8-FITC clone RPA-T8, CD33-PE clone WM53, CD14-PerCP-Cy5.5 clone HCD14, CD45RA-PE-Cy7 clone HI100, CD56-allophycocyanin clone QA17A16, CD4-Alexa Fluor 700 clone SK3; from Thermo Scientific: CD14-Alexa 700 clone Tuk4, CD3-PE-Cy5.5 clone 7D6, CD4-PE clone RPA-T4, CD19-PacBlue clone SJ25C1, CD16-PE-TexasRed clone 3G8.

The following primary antibodies were used for immunochemistry: anti-human CD31 clone JC/70A (Biocare Medical, Pacheco, CA, USA), anti-mouse CD31 clone D8V9E (Cell Signaling Technology, Danvers, MA, USA), ACE2 clone EPR34435 (Abcam, Waltham, MA, USA), polyclonal SFTPC (Seven Hills Bioreagents, Cincinnati, OH, USA), anti-human CD68 clone KP1 (LS Bio, Seattle, WA, USA), anti-human CD61 clone ARC0460 (Thermo Scientific), anti-human CD4 clone SP35 and anti-human CD8 clone SP57 (Roche, Basel, Switzerland), and anti-human CD20 clone L26 Dako Omnis (Agilent, Santa Clara, CA, USA). The secondary antibody used in this study included HRP Goat anti-Rabbit IgG (H&L) (Vector Laboratories, Burligame, CA, USA). For mouse derived primary antibodies, a linker antibody (Abcam) was used prior to application of the secondary antibody to prevent non-specific binding. DAB and purple chromogens (Roche) and chromogens used for TSA-conjugated Opal 480, 520, 570, 620, and 690 fluorophores (Akoya Biosciences, Marlborough, MA, USA) were utilized to develop immunohistochemical assays. The following anti-SARS-CoV-2 antibodies were used for immunohistochemistry: rabbit polyclonal anti-SARS-CoV Nucleoprotein (Novus Biological, Littleton, CO, USA), mouse monoclonal anti-SARS-CoV-2 Spike clone 2B3E5 (This antibody was used in this study as clone E7U60, which was the pre-production clone ID of clone 2B3E5; Cell Signaling Technology).

### Animal work

#### Institutional approvals

All animal experiments described in this study were performed in accordance with protocols (number 1930) that were reviewed and approved by the Institutional Animal Care and Use and Committee of Princeton University (#1930) and Boston University (PROTO202000020). All mice were maintained in facilities accredited by the Association for the Assessment and Accreditation of Laboratory Animal Care (AAALAC). All replication-competent SARS-CoV-2 experiments were performed in a biosafety level 3 laboratory (BSL-3) at the Boston University National Emerging Infectious Diseases Laboratories (NEIDL).

#### Mouse models and housing

NOD *Rag1^-/-^ IL2Rg^null^* mice (NOD.Cg-*Rag1^tm1Mom^Il2rg^tm1Wjl^*^/SzJ^, were obtained from the Jackson Laboratory, catalog number 007799). NRG-Flk2-/- (NRGF) mice (NOD.Cg-*Rag1^tm1Mom^ Flt3^tm1Irl^ Il2rg^tm1Wjl^*/J) were generated as described previously (Douam et al., 2018) and are available at The Jackson Laboratory (Bar Harbor, ME, USA) (catalog number 033127). NRG and NRGF mice were maintained at the Laboratory Animal Resource Center at Princeton University prior to engraftment with human tissues and shipment to the NEIDL. Heterozygous K18-hACE2 C57BL/6J mice of both sexes (strain 034860, 2B6.Cg-Tg(K18-ACE2)2Prlmn/J) were obtained from The Jackson Laboratory and maintained at the NEIDL at Boston University. In the NEIDL BSL-3 facility, mice were group-housed by sex in Tecniplast green line individually ventilated cages (Tecniplast, Buguggiate, Italy). Mice were maintained on a 12:12 light cycle at 30-70% humidity and provided sulfatrim-containing (NRG/NRGF) or ad-libitum water (K18-hACE2) and standard chow diets (LabDiet, St. Louis, MO, USA).

#### Generation of mice engrafted with human fetal lung xenografts (fLX)

Fetal lung tissues within a gestational age range of 18 to 22 weeks were obtained from Advanced Biosciences Resources (Alameda, CA, USA). Upon receipt, fetal lung tissue was trimmed of visible connective tissue before the lung was processed into cubes (3-5 mm/side) and placed into DMEM. NRG and NRGF mice (greater than 6 weeks of age) were anesthetized using isoflurane and placed in prone position. A midline incision was made along the skin of the upper back of the mouse. Forceps were used to create subdermal pockets on either side of the midline incision. A piece of lung was dried on a sterile drape and coated with Corning (Corning, NY, USA) Matrigel Basement Membrane Matrix (product number 354234). Matrigel-coated lung pieces were inserted into each of the subdermal pockets. Skin clips were used to secure the incision. Mice were used for infections 10-15 weeks following engraftment. Fetal lung xenografts derived from eight different donors were used in this study.

#### Isolation of human CD34+ hematopoietic stem cell (HSC)

Human fetal livers (16-22 weeks of gestational age) were procured from Advanced Bioscience Resources. Fetal liver was homogenized and incubated in digestion medium (HBSS with 0.1% collagenase IV (Sigma-Aldrich, Darmstadt, Germany), 40 mM HEPES, 2 M CaCl2 and 2 UU/ml DNAse I (Roche) for 30 min at 37°C. Human CD34+ HSC were isolated using a CD34+ HSC isolation kit (Stem Cell Technologies, Cambridge, MA, USA) according to the manufacturer’s protocol. Purification of human CD34+ cells were assessed by flow cytometry using an anti-human CD34-FITC antibody (clone 581, BD Biosciences). Fetal liver derived from three different donors were used in this study.

#### Generation of human immune system-engrafted mice

3-5 weeks post fLX engraftment, NRGF-L mice were irradiated with 300 cGy and 7-10x10^5^ human CD34+ HSC were injected intravenously 4-6 h after irradiation. Male and female mice transplanted with CD34+ HSC derived from three different human donors were used in this study. Twelve weeks post HSC engraftment, peripheral levels of humanization were checked. Mice with peripheral engraftment level >40% were enrolled in the study. One-week prior SARS-CoV-2 infection, NRGF-L mice were injected intravenously (tail vein) with 2×10^11^ copies of AAV-Flt3LG resuspended in 200 µl of 1X phosphate-buffered saline (PBS) containing 35nM NaCL, 0.002%pluronic F-68 and 5%glycerol.

#### Inoculation of humanized mice by intra-fetal lung xenograft injection with SARS-CoV-2

Ten to fifteen weeks post engraftment, NRG-L and HNFL mice of both sexes were inoculated via intra-fetal lung xenograft (intra-fLX) injection with 10^4^ or 10^6^ PFU of SARS-CoV-2 in 50 µL of sterile 1X PBS. Inoculations were performed under 1-3% isoflurane anesthesia. Either one or both implants were inoculated by direct injection into the fLX. Animals were euthanized at day two or day seven post inoculation.

#### Intranasal inoculation of K18-hACE2 with SARS-CoV-2

10-12 week old K18-hACE2 mice of both sexes were intranasally inoculated with 10^6^ PFU of SARS-CoV-2 in 50 µl of sterile 1X PBS. Inoculations were performed under 1-3% isoflurane anesthesia. Animals were euthanized either prior to infection, at 2DPI, at 7DPI, or when they reached euthanasia criteria (as defined in related IACUC protocol).

#### Tissue collection and lung inflation for histology

At the indicated endpoints, mice were anesthetized using 1-3% isoflurane, followed by euthanasia using an overdose of ketamine/xylazine. For lung inflation specifically, a midline incision was made to open the abdominal cavity of the mouse. After the diaphragm was punctured, the rib cage was cut from sternum to chin to open the thoracic cavity. Following opening of the rib cage, skin and subcutaneous tissue were removed to expose the trachea, and an 18G catheter was inserted into the trachea and secured with a ligature. The lungs were insufflated with 1.5 mL of low-melt agarose and removed from the mouse after agarose solidification, then placed in 10% NBF for fixation for a minimum of 72 hours.

### Virus

#### SARS-CoV-2 isolate stock

All replication-competent SARS-CoV-2 experiments were performed in a BSL-3 facility at the Boston University National Emerging Infectious Diseases Laboratories. The clinical isolate named 2019-nCoV/USA-WA1/2020 strain (NCBI accession number: MN985325) of SARS-CoV-2 was obtained from BEI Resources (Manassas, VA, USA). To generate the passage 1 (P1) virus stock, Vero E6 cells, pre-seeded the day before at a density of 10 million cells, were infected in T175 flasks with the master stock, diluted in 10 ml final volume of Opti-MEM (ThermoFisher Scientific, Waltham, MA, USA). Following virus adsorption to the cells at 37°C for 1 h, 15 ml DMEM containing 10% FBS and 1X penicillin/streptomycin was added to the flask. The next day, media was removed, the cell monolayer was rinsed with 1X PBS, pH 7.5 (ThermoFisher Scientific) and 25 ml of fresh DMEM containing 2% FBS was added. Two days later, when the cytopathic effect of the virus was clearly visible, culture medium was collected, filtered through a 0.22 µm filter, and stored at −80°C. Our P2 working stock of the virus was prepared by infecting Vero E6 cells with the P1 stock, at a multiplicity of infection (MOI) of 0.1. Cell culture media was harvested at 2DPI and 3DPI, and after the last harvest, ultracentrifuged (Beckman Coulter Optima L-100k; SW32 Ti rotor) for 2 h at 25,000 rpm (80,000 x *g*) over a 20% sucrose cushion (Sigma-Aldrich). Following centrifugation, the media and sucrose were discarded, and pellets were left to dry for 5 min at room temperature. Pellets were then resuspended overnight at 4°C in 500 µl of 1X PBS. The next day, concentrated virions were aliquoted and stored at −80°C.

#### Production of recombinant SARS-CoV-2 expressing NanoLuc Luciferase

A recombinant SARS-CoV-2 expressing NanoLuc Luciferase (rSARS-CoV-2 NL) (Xie et al., 2020) was graciously provided by the Laboratory of Pei-Yong Shei. To propagate the virus, a day prior to propagation 10 million Vero E6 cells were seeded in a T-175 flask, 10 µL of rSARS-CoV-2 NL virus stock was diluted in 10 mL of OptiMEM. Virus was incubated on cells for 1 h at 37°C then 15 mL of DMEM containing 10% FBS and 1% penicillin/streptomycin was added. The next morning, media was removed, cells were washed with 1X PBS and 25 mL of fresh DMEM containing 2% FBS was added. Virus was incubated for an additional 48 h, supernatant was collected, filtered through a 0.22 µm filter, and stored at −80°C. Viral stock was thawed and concentrated by ultracentrifugation (Beckman Coulter Optima L-100k; SW32 Ti rotor) at 25,000 x *g* for 2 h at 4°C on a 20% sucrose cushion (Sigma-Aldrich, St. Louis, MO). Media and sucrose were discarded, pellets were dried for 5 min at room temperature, then viral pellets were suspended in 100 µL of 1X PBS at 4°C overnight. On the next day, concentrated virus was aliquoted and stored at −80°C.

#### SARS-CoV-2 titering

The titer of our viral stocks was determined by plaque assay. Vero E6 cells were seeded into a 12-well plate at a density of 2.5×10^5^ cells per well and infected the next day with serial 10-fold dilutions of the virus stock for 1 h at 37°C. Following virus adsorption, each well was supplemented with 1 ml of overlay media, consisting of 2X DMEM supplemented with 4% FBS and mixed at a 1:1 ratio with 1.2% Avicel (DuPont, Wilmington, DE, USA; RC-581). Three days later, the overlay media was removed, the cell monolayer was washed with 1X PBS and fixed for 1 h at room temperature with 10% neutral buffered formalin (ThermoFisher Scientific). Following formalin removal, fixed cells were then washed with 1X PBS and stained for 1 h at room temperature with 0.1% crystal violet (Sigma-Aldrich) prepared in 10% ethanol/water. After rinsing with tap water, the number of plaques was counted and the virus titer was calculated.

#### Generation of AAV-Flt3LG

The pAB269 AAV backbone containing AAV2 ITRs was kindly provided by Markus Grompe (OHSU, Oregon, USA). The plasmid was digested with PacI/MluI HF. The FLT3 was PCR amplified from pAL119-FLT3L (Addgene, item #21910), and the TBG was amplified from pX602-AAV-TBG:NLS-SaCas9-NLS-HA-OLLAS-bGHpA;U6::BsaI-sgRNA (Addgene, item #61593). The FLT3, TBG, and BGH PCR products had 15 bp of overlapping sequence with adjacent inserts and backbone and were assembled with In-Fusion (Takara Bio, Mountain View, CA, USA) to create the final construct. AAV-293 cells (Agilent) at 50% confluency in 15 cm dishes were transfected via the calcium phosphate method with 22.5 µg XR8 (NGVB, Indianapolis, IN), 7.5 µg pHelper (Agilent), 7.5 µg of pAB269-TBG-FLT3 LG-BGH per plate. Media was collected every 24 h for 72 h total. After 72 h, the media was treated with a 5X solution of 40% PEG8000 and 2.5 M NaCl to precipitate the AAV for 2 h at 4°C before being spun down at 4300 x *g* for 20 min. Cells from plates were scraped, washed with PBS, and resuspended in hypotonic buffer (10 mM HEPES, 1.5 mM MgCl2, 10 mM KCl, 0.35 mg/ml spermine) on ice for 10 min before 1 ml restore buffer (62.5% sucrose wt/vol in hypotonic buffer) was added. Cell membranes were sheared in a 15 ml Kontes dounce homogenizer and nuclei were spun down at 500 x *g* for 10 min. AAV from PEG precipitate was resuspended in 6 ml high salt buffer (2.5 mM KCl, 1 mM MgCl2, 1 M NaCl in PBS) and added to nuclei that had been resuspended in 1 ml low salt buffer (2.5 mM KCl, 1 mM MgCl2, in PBS). Lysate was treated with 250 units of Benzonase (Sigma) at 37°C for 30 min and then spun at 4300 x *g* for 30 min before being loaded onto an iodixanol gradient. AAV was spun at 38,000 rpm in an SW41 rotor for 3 h at 16°C. AAV was collected from the 40% iodixanol layer and buffer was exchanged to AAV storage buffer (PBS with 35 mM NaCl, 0.002% pluronic F-68, 5% glycerol) in a 100 MWCO centrifugal filter column (MilliporeSigma, Burlington, MA, USA). Samples were analyzed via silver stain to check purity and qPCR to quantify.

### Tissue processing

#### Single cell suspension from whole blood

Blood (200 µl) was collected through submandibular bleeding and transferred into EDTA capillary collection tubes (Microvette 600 K3E; Sarstedt, Nümbrecht, Germany). Cells were separated from serum through centrifugation, and red blood cells were lysed with 1X lysis buffer (BD Pharm Lyse, BD Biosciences) for 15 min at room temperature in the dark. Following lysis and quenching with 10% (v/v) FBS DMEM media, blood cells were then washed twice with a 1% (v/v) FBS-PBS solution (FACS Buffer) before antibody staining.

#### Single cell suspension from fLX

Fetal lung xenografts were collected and placed in Roswell Park Memorial Institute Medium (RPMI) with 10% FBS. To generate single cell suspensions, lung tissues were placed on a 60 mm dish and minced using a disposable scalpel. Tissue pieces were transferred to a 15 mL conical tube with 3 mL of digestion buffer (HBSS minus Ca^2+^, Mg^2+^, and phenol red, 0.5 mg/mL Liberase TM, 1 mg/mL DNase I) and incubated at 37°C for 30 min with agitation every 10 min. Minced pieces were transferred to a 70 µm strainer on a 50 mL tube and mashed through using the plunger of a 3 mL syringe plunger. The strainer was washed two times with 1 mL of FACS buffer (1X PBS with 1% (v/v) FBS) and the cell suspension was centrifuged at 300 x *g* for 5 min at 4°C. The cell pellet was resuspended in 1 mL of ACK lysing buffer (ThermoFisher Scientific; #A1049201) and incubated for 2 min at room temperature. After incubation, 9 mL of FACS buffer was added to quench the lysis, samples were centrifuged at 300 x *g* for 5 min at 4°C, and the cell pellet was resuspended in 1 mL of FACS buffer prior to antibody staining.

#### Single cell suspension from spleen

Spleen was collected and placed in RPMI with 10% FBS. To generate single cell suspensions, a 70 µm strainer was placed into one well of a 6-well plate with 4 mL of FACS buffer. Whole spleen was then placed onto the strainer and mashed through the strainer using a 3 mL syringe plunger. After the strainer was washed twice with 1 mL of FACS buffer, the resultant single cell suspension was transferred to a 15 mL conical tube and samples were centrifuged at 300 x *g* for 5 min at 4°C. The cell pellet was resuspended in 1 mL of ACK lysing buffer and incubated for 2 min at room temperature. After incubation, 9 mL of FACS buffer was added to quench the lysis, samples were centrifuged at 300 x *g* for 5 min at 4°C, and the cell pellet was resuspended in 1 mL of FACS buffer.

### RNA extraction

#### Generation of cell lysates for total RNA extractions

Tissues were collected from mice and placed in 600 µL of RNAlater (MilliporeSigma: #R0901500ML) and stored at −80°C. For processing, 20–30 mg of tissue was taken and placed into a 2 mL tube with 600 µL of RLT buffer with 1% β–mercaptoethanol and a 5 mm stainless steel bead (Qiagen, Hilden, Germany: #69989). Tissues were then dissociated using a Qiagen TissueLyser II (Qiagen) with the following cycle: two min dissociation at 1800 oscillations/min, one min rest, two min dissociation at 180 oscillations/min. Samples were then subject to centrifugation at 13,000 rpm for 10 min at room temperature and supernatant was transferred to a new 1.5 mL tube. RNA extractions were performed using a Qiagen RNeasy Plus Mini Kit (Qiagen: #74134), according to the manufacturer’s instructions, with an additional on-column DNase treatment (Qiagen: #79256). RNA was eluted in 30 µL of RNase/DNase free water.

#### RNA extraction from serum

Viral RNA was extracted from serum using a Zymo Viral RNA extraction kit (Zymo Research, Irvine, CA, USA: #R1035) following the manufacturers protocol. Briefly, serum was mixed with RNA/DNA shield (Zymo) at a 1:1 ratio. RNA buffer was then added to the serum (2:1 ratio) and passed through a column by centrifugation at 13,000 x *g*. The column was then washed twice, and RNA was eluted with 15 µL of RNase/DNase free water.

### Flow cytometry

For all flow cytometry experiments, flowcytometric analysis was performed using an LSRII Flow Cytometer (BD Biosciences). Flow cytometry fluorophore compensation for antibodies was performed using an AbC™ Anti-Mouse Bead Kit (ThermoFisher Scientific). Flow cytometry data were analyzed using FlowJo software (TreeStar, Ashland, OR, USA).

#### Quantification of peripheral human chimerism in HNFL mice

2-4×10^6^ PBMCs of human or murine origin were isolated as described above and stained for 1 h at 4°C in the dark with an antibody cocktail targeting human(h)CD45, mouse CD45, hCD3, hCD4, hCD8, hCD16, hCD19, hCD11c, hCD56 and hCD14. Following washing with FACS Buffer, cells were fixed with fixation buffer (1% (v/v) FBS, 4% (w/v) PFA in PBS) for 30 min at 4°C in the dark. Chimerism of all humanized mice was assessed by quantifying the following human populations: Human CD45^+^, human CD45^+^ murine CD45^-^; T-cells, CD45^+^ CD3^+^; CD4^+^ T cells, CD45^+^ CD3^+^ CD4^+^; CD8^+^ T cells, CD45^+^ CD3^+^ CD8^+^; CD45^+^ CD16^+^ leukocytes; B-cells, CD45^+^ CD19^+^; conventional dendritic cells, CD45^+^ CD11c^+^; NK/NKT cells, CD45^+^ CD56^+^; Monocytes, CD45^+^ CD14^+^.

#### Antibody staining and flow cytometry analysis of NRG-L splenocytes and fLX

2-4×10^6^ splenocytes or fetal lung cells of human or murine origins were isolated as described above and stained for 1 h at 4°C in the dark with an antibody cocktail targeting mouse CD45, human(h)CD45, hCD19, hCD3, hCD33, hCD11c, hCD56, hCD68, hCD123, hCD14. Following washing with FACS Buffer, cells were fixed with fixation buffer for 30 min at 4°C in the dark. Human immune cell subsets from naïve NRG-L fLX were gated as follows: Human CD45^+^, human CD45^+^ murine CD45^-^; T-cells, CD45^+^ CD3^+^; CD11c/CD33, CD45^+^ CD3^-^ CD19^-^ CD33^+^ CD11c^+^; CD56, CD45^+^ CD3^-^ CD19^-^ CD56^+^.

#### Antibody staining and flow-cytometry analysis of HFNL fLX

After generation of single cell suspension, 5×10^5^ − 1×10^6^ cells were used for flow cytometry staining. Cells were centrifuged at 300 X *g* for 5 min at 4°C. The cell pellet was resuspended in a mix of 22.5 µL FACS buffer and 2.5 µL of FcX (Biolegend; #422302) and incubated for 10 min at room temperature. After blocking, 25 µL of antibody cocktail targeting hCD3, hCD20, hCD16, hHLA-DR, hCD45, hCD8, hCD4, hCD33, hCD45RA, hCD56, hCD14, mCD45, and containing a LIVE/DEAD viability dye (ThermoFisher Scientific) was added to each sample and incubated in the dark for 30 min at 4°C. After staining, 1 mL of FACS buffer was added to each sample, samples were centrifuged at 300 x *g* for 5 min, washed with 1 mL FACS buffer, centrifuged at 300 x *g* for 5 min, and then fixed in 200 µL 4% PFA for 30 min. After fixation cells were washed twice with 1 mL FACS buffer, resuspended in FACS buffer, and stored protected from light at 4°C until analysis. Human immune cell subsets were gated as follows: human CD45^+^, hCD45^+^ mCD45^-^; human CD3^+^, hCD45^+^ hCD3^+^; human CD4^+^, hCD45^+^ hCD3^+^ hCD4^+^; human CD8^+^, hCD45^+^ hCD3^+^ hCD8^+^; CD20^+^, hCD45^+^ hCD3^-^ hCD20^+^; human CD56^+^, hCD45^+^ hCD3^-^ hCD20^-^ hCD33^-^ hCD56+.^+^.

### Viral quantification

#### SARS-CoV-2 RT-qPCR

To determine SARS-CoV-2 RNA copies, RT-qPCR for SARS-CoV-2 E protein was performed using a One-Step Taqman-based system. Briefly, a 20 µL reaction mixture containing 10 µL of Quanta qScript™ XLT One-Step RT-qPCR ToughMix® (VWR, Radnor, PA, USA; #76047-082), 0.5 µM Primer E_Sarbeco_F1 (ACAGGTACGTTAATAGTTAATAGCGT), 0.5 µM Primer E_Sarbeco_R2 (ATATTGCAGCAGTACGCACACA), 0.25 µM Probe E_Sarbeco_P1 (FAM-ACACTAGCCATCCTTACTGCGCTTCG-BHQ1), and 2 µL of total RNA was subjected to One-Step RT-qPCR using Applied Biosystems QuantStudio 3 (ThermoFisher Scientific), with the following cycling conditions; reverse transcription for 10 min at 55°C and denaturation at 94°C for 3 min followed by 45 cycles of denaturation at 94°C for 15 sec and annealing/extension at 58°C for 30 sec. Ct values were determined using QuantStudio^TM^ Design and Analysis software V1.5.1. Calculations for RNA copies/mL were determined using a SARS-CoV-2 E RNA as a standard.

#### Quantification of infectious particles by plaque assay

SARS-CoV-2 infectious particles in infected fLX were quantified by plaque assay. After euthanizing mice, tissues were collected in 500 µL of RNAlater (MilliporeSigma: # R0901500ML) and stored at −80°C. The day prior to experiments, 8×10^4^ cells per well were seeded in a 24-well plate. Between 20 and 30 mg of tissue was placed into 500 µL of OptiMEM (ThermoFisher Scientific). Tissues were homogenized using a Qiagen TissueLyser II (Qiagen) by two dissociation cycles (two min at 1800 oscillations/min) with one min rest in between. Samples were then subjected to centrifugation at 13,000 rpm for 10 min at room temperature, and supernatant was transferred to a new 1.5 mL tube. From this, 1:10^2^ – 1:10^7^ dilutions were made in OptiMEM and 200 µL of each dilution were plated onto a 24-well plate. Supernatant was incubated at 37°C for 1 h with gentle rocking of the plate every 10 min. After viral adsorption, 1 mL a 1:1 mixture of 2X DMEM 4% FBS and 2.4% Avicel (Dupont) was overlaid into each well. Plates were incubated for 72 h at 37°C with 5% CO_2_. Avicel was then removed, cells were washed twice with 1X PBS, and then cells were fixed in 10% buffered formalin (ThermoFisher Scientific) for 1 h. After fixation, formalin was removed, and cells were stained with 0.1% crystalline violet in 10% ethanol/water for 1 h and washed with tap water. Number of plaques were counted, and infectious particles (PFU/mg of tissue) were calculated.

### Histology and microscopy

#### Histologic processing and analysis

Tissue samples were fixed for 72 h in 10% neutral buffered formalin, processed in a Tissue-Tek VIP-5 automated vacuum infiltration processor (Sakura Finetek USA, Torrance, CA, USA), followed by paraffin embedding using a HistoCore Arcadia paraffin embedding station (Leica, Wetzlar, Germany). Generated formalin-fixed, paraffin-embedded (FFPE) blocks were sectioned to 5 μm using a RM2255 rotary microtome (Leica), transferred to positively charged slides, deparaffinized in xylene, and dehydrated in graded ethanol. Tissue sections were stained with hematoxylin and eosin for histologic examination, while unstained slides were used for immunohistochemistry. Qualitative and semi-quantitative histomorphological analyses were performed by a single board-certified veterinary pathologist (N.A.C.). An ordinal scoring system was developed to capture the heterogeneity of histologic findings in fLXs. Individual histologic findings that were scored included: syncytial cells, hyaline membrane, intra-airspace neutrophils and necrosis, hemorrhage, edema, denuded pneumocytes, capillary fibrin thrombi, intermediate vessel fibrin thrombi and coagulative necrosis. The entire fLX was examined at 200x with a DM2500 light microscope (Leica) using the following criteria: 0 = not present, 1 = found in <5% of fields, 2 = found in >5% but <25% of fields, or 3 = found in >25% of fields. Several criteria were also restricted to ‘not observed’ (0) or ‘observed’ (1). Scores were combined to generate a cumulative lung injury score.

#### Multispectral fluorescent imaging

Fluorescently labeled slides were imaged using either a Mantra 2.0^TM^or Vectra Polaris ^TM^ Quantitative Pathology Imaging System (Akoya Biosciences). To maximize signal-to-noise ratios, fluorescently acquired images were spectrally unmixed using a synthetic library specific for the Opal fluorophores used in each assay plus DAPI. An unstained fLX section was used to create an autofluorescence signature that was subsequently removed from multispectral images using InForm software version 2.4.8 (Akoya Biosciences).

#### Image analysis of multiplex immunohistochemistry

Digitized whole slide scans were analyzed using the image analysis software HALO v3.2 (Indica Labs, Inc., Corrales, NM, USA). Slides were manually annotated to select regions of interest, excluding accessory skin and adipose tissue, to ensure inclusion of only the fLX. Visualization thresholds were adjusted in viewer settings to minimize background signal identification and maximize specificity of signals for each sample. Quantitative positive pixel analysis outputs were obtained using the Area Quantification (AQ) module, which reports total area of immunoreactivity of a specified parameter within a region of annotated interest. Values are given as a percentage of total tissue area analyzed. Minimum dye intensity thresholds were established using the real-time tuning field of view module to accurately detect positive immunoreactivity. For quantitative cellular phenotyping, the fluorescence Highplex (HP) module was utilized. Cells are identified using DAPI to segment individual nuclei. Minimum cytoplasm and membrane thresholds are set for each dye to detect positive staining within a cell. Parameters are set using the real-time tuning mechanism that was tailored for each individual biopsy based on signal intensity. Phenotypes are determined by selecting inclusion and exclusion parameters relating to stains of interest. This algorithm produces a quantitative output for each cell phenotype standardized to the area analyzed (cells/µm^2^).

#### Transmission electron microscopy

Tissue samples were fixed for 72 h in a mixture of 2.5% glutaraldehyde and 2% formaldehyde in 0.1 M sodium cacodylate buffer (pH 7.4). Samples were then washed in 0.1 M cacodylate buffer and postfixed with 1% osmiumtetroxide (OsO4)/1.5% potassiumferrocyanide (KFeCN6) for 1 h at room temperature. After washes in water and 50 mM maleate buffer pH 5.15 (MB), the samples were incubated in 1% uranyl acetate in MB for 1 h, washed in MB and water, and dehydrated in grades of alcohol (10 min each: 50%, 70%, 90%, 2×10 min 100%). The tissue samples were then put in propyleneoxide for 1 h and infiltrated overnight in a 1:1 mixture of propyleneoxide and TAAB Epon. The following day the samples were embedded in fresh TAAB Epon and polymerized at 60°C for 48 h. Semi-thin (0.5 μm) and ultrathin sections (50-80 nm) were cut on a Reichert Ultracut-S microtome (Leica). Semi-thin sections were picked up on glass slides and stained with toluidine blue for examination by light microscopy to find affected areas in the tissue. Ultrathin sections from those areas were picked up onto formvar/carbon coated copper grids, stained with 0.2% lead citrate and examined in a JEOL 1200EX transmission electron microscope (JOEL, Akishima, Tokyo, Japan). Images were recorded with an AMT 2k CCD camera.

### Proteomics and transcriptomic analysis

#### Mass spectrometry sample preparation

Inactivated protein extracts were sonicated with a Branson probe sonicator and were then quantified via Bradford assay. The samples were diluted with 100 mM Tris, pH 8.5 buffer to lower the GuHCl concentration to 0.75 M. Lysate proteins were then digested by adding trypsin (Pierce, Waltham, MA, USA) at a 1:50 ratio (enzyme:protein, w/w) and incubating the samples overnight at 37°C with shaking. Trypsin digestion was terminated with the addition of TFA to below pH 3 and the peptide digests were desalted via reversed-phase C18 columns (Sep-Pak, Waters, Mildford, MA, USA) with a wash buffer of 0.1% TFA and elution buffer of 60% acetonitrile. The desalted peptides were then quantified with a Quantitative Colorimetric Peptide Assay (Pierce). Each sample, comprising 100 μg peptides, was TMT-labeled with TMTPro 16plex reagents (ThermoFisher Scientific: # A44520) as per the manufacturer’s protocol. Labeled peptides were again desalted on a C18 column prior to basic reversed-phase fractionation.

TMT-labeled peptides were fractionated via basic reversed-phase chromatography on the Agilent 1200 series HPLC instrument equipped with the XBridge Peptide BEH C18 column (130A°, 3.5 mm, 4.6 mm X 250 mm, Waters Corporation). Prior to loading peptides, the C18 column was washed with 100% methanol and equilibrated with Buffer A (0.1% ammonium hydroxide and 2% acetonitrile). Peptides were injected via the autosampler and eluted from the column using a gradient of mobile phase A (2% ACN, 0.1% NH4OH) to mobile phase B (98% ACN, 0.1% NH4OH) over 48 min at a flowrate of 0.4 mL/min. The 48 fractions collected were orthogonally concatenated into 12 pooled fractions. Three percent of each fraction was aliquoted and saved for global proteomic profiling and the remaining 97% of peptides were used for phosphopeptide enrichment using Fe-NTA magnetic beads (CubeBiotech, Monheim am Rhein, Germany) (Leutert et al., 2019). Briefly, the fractionated peptides were dried and resuspended in binding buffer (80% Acetonitrile and 01% TFA). Before being added to the peptides, the Fe-NTA beads were washed with binding buffer. Peptides were then incubated with the Fe-NTA bead slurry for 20 min in a Kingfisher Apex magnetic bead transferring system (ThermoFisher Scientific) before being moved into wash wells. Beads with bound phosphopeptides were washed twice in binding buffer, after which phosphopeptides were serially eluted twice by moving the beads into wells containing 200 µL of elution buffer (50% acetonitrile and 2.5% ammonium hydroxide). Both phosphopeptide eluates corresponding to an orthogonal fraction were combined prior to drying in a speedvac.

#### Mass spectrometry analysis

Multiplexed peptide fractions from each time point were resuspended in mobile phase A solvent (2% acetonitrile and 0.1% formic acid) to be analyzed on the Exploris 480 mass spectrometer equipped with FAIMS (ThermoFisher Scientific). The mass spectrometer was interfaced to the Easy nanoLC1200 HPLC system (ThermoFisher Scientific). Briefly, the peptides were first loaded onto a reverse-phase nanotrap column in mobile phase A, (75 mm i.d. 3 2 cm, Acclaim PepMap100 C18 3 mm, 100 A°, ThermoFisher Scientific) and separated over an EASY-Spray column, (ES803A, ThermoFisher Scientific) using a gradient (6% to 19% over 58 min, then 19% to 36% over 34 min) of mobile phase B (0.1% formic acid, 80% acetonitrile) at a flowrate of 250 nL/min. The mass spectrometer was operated in positive ion mode with a capillary temperature of 275°C and a spray voltage of 2500 V. All data was acquired with the mass spectrometer operating in data dependent acquisition (DDA) mode, with FAIMS cycling through one of three compensation voltages (-50V, -57V, -64V) at each full scan. Precursor scans were acquired at a resolution of 120,000 FWHM with a maximum injection time of 120 ms in the Orbitrap analyzer. The following 0.8 sec were dedicated to fragmenting the most abundant ions at the same FAIMS compensation voltage, with charge states between 2 and 5, via HCD (NCE 33%) before analysis at a resolution of 45,000 FWHM with a maximum injection time of 60 ms. Phosphopeptides were analyzed in the same manner, save for the injection time being raised to 150 ms to allow for lower abundant analyte to fill the trap.

#### Analysis of raw mass spectrometry data

All acquired MS/MS spectra were simultaneously searched against the complete SwissProt human proteome (downloaded on 2020-10-20), the complete SwissProt mouse proteome (downloaded on 2020-10-20), and the Uniprot SARS-CoV-2 proteome (For side-by-side NRG-L vs. HNFL experiment only; downloaded on 2020-05-03) using MaxQuant (Version 1.6.7.0), which integrates the Andromeda search engine. TMT reporter ion quantification was performed using MaxQuant with default settings. Briefly, enzyme specificity was set to trypsin and up to two missed cleavages were allowed. Cysteine carbamidomethylation was specified as fixed modification whereas oxidation of methionine and N-terminal protein acetylation were set as variable modifications. For phosphopeptides serine, threonine and tyrosine, phosphorylation were specified as variable modifications. Precursor ions were searched with a maximum mass deviation of 4.5 ppm and fragment ions with a maximum mass deviation of 20 ppm. Peptide and protein identifications were filtered at 1% FDR using the target-decoy database search strategy (Elias and Gygi, 2007). Proteins that could not be differentiated based on MS/MS spectra alone were grouped to protein groups (default MaxQuant settings). A threshold Andromeda score of 40 and a threshold delta score of 8 was applied to phosphopeptides. The MaxQuant output files designated ‘‘Phospho(STY)sites’’ and ‘‘proteinGroups’’ were filtered to remove entries that were either entirely mouse, or in the case of completely homologous peptides, had annotations of both mouse and human. These two files, filtered to contain only accessions related to human proteins, were used for data normalization and other statistical analysis using in-house generated scripts in the R environment.

#### Bulk RNA sequencing

Total RNA was processed from fLX as described above, and sent to BGI genomics (Hong Kong, China) for library preparation and sequencing (Pair-ends, 100 bp, 20M reads per sample). Raw FASTQ files were quality-checked with FastQC v0.11.7. Reads were found to be excellent quality and were aligned to the combined human (GRCh38, Ensembl 101) and mouse (GRCm38, Ensembl 101) genomes with STAR v2.7.1a followed by quantification with featureCounts v1.6.2. Quality was checked with MultiQC v1.6. All samples passed quality thresholds of >75% sequences aligned and >15 million aligned reads per sample. Significantly up- and downregulated genes were identified with DESeq2 v1.23.10 in R v3.6.0. Three treatment groups were compared to uninfected samples in turn: 2DPI, 7DPI and 7DPI contralateral. P-values were FDR-adjusted, and log_2_ fold change was shrunk with the *apeglm* method. Significance was determined by an FDR-adjusted *p*<0.01 and a shrunken log_2_ fold change > 2 or < -2. DESeq2 result were imported into Ingenuity Pathway Analysis (IPA; Service curated by Qiagen; Access provided through the Boston University Genome Science Institute) (Kramer et al., 2014), and a canonical pathway enrichment analysis was performed using the default settings and the same differential expression thresholds as before (shrunken log2 fold change > 2 or < -2 and FDR-adjusted p-value < 0.01).

#### Single cell barcoding and sequencing

Following fLX processing into single cell suspension as described above, cells were frozen down in a 90% FBS (Bio-Techne, R&D systems) 10% DMSO solution (ThermoFisher Scientific) and kept at −80°C. Four to five days following freezing, cells were thawed, and viability was assessed using Trypan blue (Fisher Scientific). Samples with ≥90% viability were then processed using the Chromium Next GEM Single Cell 3’ GEM, Library & Gel Bead Kit v3.1, as per manufacturer instructions, and single cell barcoding was performed using a Chromium instrument (10x genomics) located in the NEIDL BSL-3. Reverse transcription of RNAs was performed in the BSL-3 using a thermocycler (Applied Biosciences), and cDNA was then removed from containment. Full-length, barcoded cDNAs were then amplified by PCR to generate sufficient mass for library construction. Enzyme fragmentation, A tailing, adaptor ligation and PCR were then performed at the Boston University single-cell sequencing core to obtain final libraries containing P5 and P7 primers used in Illumina bridge amplification. Size distribution and molarity of resulting cDNA libraries were assessed via Bioanalyzer High Sensitivity DNA Assay (Agilent Technologies, USA). All cDNA libraries were sequenced on an Illumina NextSeq 500 instrument at the Boston University microarray and sequencing core according to Illumina and 10x Genomics guidelines with 1.4-1.8pM input and 1% PhiX control library spike-in (Illumina, USA).

#### Preprocessing and quality control of single-cell data

The 10X CellRanger tool was used for demultiplexing, alignment, identification of cells, and counting of unique molecular indices (UMIs). Specifically, the CellRanger *makefastq* command was used to demultiplex raw base call (BCL) files generated by Illumina sequencers into FASTQ files. The CellRanger *count* command was used to perform alignment and create UMI count matrices using parameters --expect-cells=6000. Multiple sequence alignments were performed. The reads were first aligned to the combined GRCh38 & mm10 reference. DecontX (Yang et al., 2020b) was then applied to estimate the cell contamination scores. The reads were then aligned to the GRCh38 and mm10 references individually, separating the mouse and human cells. Cells with more than 350 human counts and less than 250 mouse counts were classified as human, and vice versa. Finally, the reads were also aligned to a custom reference constructed by adding the SARS-CoV-2 transcriptome to the GRCh38 reference genome. Droplets with at least 500 UMIs underwent further quality control with the SCTK-QC pipeline [REF]. The median number of UMIs was 2,685, the median number of genes detected was 934, the median percentage of mitochondrial reads was 0%, and the median contamination estimated by decontX (Yang et al., 2020b) was 2% across cells from all samples. Cells with less than 3 counts, less than 3 genes detected were excluded leaving a total of 42,182 cells for the downstream analysis.

#### Clustering of single-cell data with Celda

The *celda* package was used to bi-clustering genes into modules and cells into subpopulations (Wang et al., 2020). The 5,000 most variable features were selected using the *seuratFindHVG* function from the *singleCellTK* package after excluding features with less than 3 counts in 3 cells. The *recursiveSplitModule* and *recursiveSplitCell* functions were used to select the model with 150 modules and 15 cell subpopulations after examining the Rate of Perplexity Change (RPC). Cells were embedded in two dimensions with UMAP using the *celdaUmap* function. Heatmaps for specific modules were generated using the *moduleHeatmap* function. Markers for each cluster were identified with the *findMarkerDiffExp* function from the *singleCellTK* package using the parameters MAST algorithm (Finak et al., 2015) and parameters FDR threshold = 0.05. Clusters were annotated manually based on the level of expression of cluster defining genes and can be found in **Supplemental Item 1**.*In vivo* imaging and data analysis

NRG-L mice were infected with 10^6^ PFU of rSARS-CoV-2 NL virus via direct fLX inoculation. For imaging, mice were injected with 5 µg (0.25 mg/kg) furimazine substrate (MedChemExpress, Monmouth Junction, NJ, USA) via tail-vein intravenous injection. Mice were then imaged using a 3D-imaging mirror gantry isolation chamber (InVivo Analytics, New York, NY, USA) and an IVIS Spectrum imager (PerkinElmer, Waltham, MA, USA). Briefly, mice were anesthetized using isoflurane (2.5%), placed into a body conforming animal mold (BCAM) (InVivo Analytics) and imaged within 5 min of injection. Mice were imaged using a sequence imaging file as follows: 60 sec open filter, 240 sec 600 nM, 60 sec open, 240 sec 620 nm, 60 sec open, 240 sec 640 nm, 60 sec open, 240 sec 660 nm, 60 sec open, 240 sec 680 nm, 60 sec open using an IVIS Spectrum imager (PerkinElmer). Data analysis of planar imaging was conducted using LivingImage (PerkinElmer). 3D reconstitution of bioluminescence signals was conducted manually by InVivo Analytics.

### Statistical Analysis

For histopathological score and viral loads/titers comparison, Kruskal-Wallis or a two-way ANOVA with Benjamini, Krieger, and Yekutieli correction for multiple comparisons were applied given the non-continuous nature of the data (i.e., viral inoculation in areas displaying differential stage of tissue development, and/or differences in fLX engraftment, and/or donor/gestational age differences). A two-way ANOVA with Benjamini, Krieger, and Yekutieli correction for multiple comparisons was also used to generate the ISG *p.value* heatmap given the non-parametric nature of these data. For quantitative analysis of hematopoietic cell infiltration (i.e. multiplex IHC and flowcytometry data), Welch’s t-test or one-way ANOVA tests were used. For comparison of viral gene count between macrophage clusters and other clusters, a Mann-Whitney t-test was used. All statistical tests and graphical depictions of results were performed using GraphPad Prism version 9.0.1 software (GraphPad Software, La Jolla, CA). For all tests, *p*≤0.05 was considered statistically significant. Statistical significance on figures and supplemental figures is labelled as follow: \**p*≤0.05, \*\**p*≤0.01, \*\*\**p*≤0.001, \*\*\*\**p*≤0.0001.

### Data deposition

The mass spectrometry proteomics data have been deposited to the ProteomeXchange Consortium via the PRIDE (Perez-Riverol et al., 2019) partner repository with the dataset identifier PXD025851. Transcriptomic data generated during this study are available through the National Center for Biotechnology Information Gene Expression Omnibus (GEO) under series accession no. GSE180063.

## SUPPLEMENTAL INFORMATION

### Supplemental Figures

**Figure S1. Related to Figure 1. Engraftment of NRG-L mice with human fetal lung xenografts (fLX).**

**A.** An NRG-L mouse engrafted with pairs of human fLX (red ellipse).

**B.** Left and right fLX following extraction from an NRG-L mouse 4 weeks post engraftment.

**C.** Bilateral fetal lung xenografts (fLX) subcutaneously implanted in NRG mice. Recruitment of subcutaneous murine blood vessels is highlighted in the inset (black arrows).

**D-E.** Representative integration of murine and human blood vessels in human fLX following engraftment. (D) Integration of murine and human blood vessels at the interface of murine subcutis (left of the black hash line) and fLX (right of the black hash line). (E) Both human and murine blood vessels are present within the fetal lung xenograft interstitium. Purple (murine CD31) and brown (human CD31) duplex IHC, 200x (top, bar=100um), 400x total magnification (bottom, bar=50um).

**F-M.** Representative fetal lung xenografts tissue sections displaying heterogenous stages of lung maturation and differentiation affiliated with differential expression of ACE2 (host receptor for SAS-CoV-2), Prosurfactant Protein C (SFTPC-Alveolar type II pneumocyte differentiation), and CD31 (endothelium-human specific). (F-G) Pseudoglandular phenotype, columnar to cuboidal pneumocytes with high expression of ACE2 and SFTPC, forming glandular like structures supported by prominent mesenchymal stroma with high vascular density. (H-I) Canalicular phenotype, low cuboidal pneumocytes with moderate sporadic expression of ACE2 and SFTPC, retraction of mesenchymal stroma, decreased vascular density, and formation of coalescing airspaces with loss of distinct glandular like structures. (J-K) Saccular phenotype, further enlargement of airspaces lined by squamous epithelium with low to moderate expression of ACE2 and SFTPC, formation of distinct septal like structures, and low vascular density. (L-M) Bronchiole differentiation, columnar to pseudostratified ciliated epithelium with apical ACE2 expression and absence of SFTPC. F,H,J,L: Hematoxylin and Eosin (H&E), magnification 100x, scale bar=200um. G,I,K,M: fluorescent multiplex immunohistochemistry (mIHC), magnification 200x, scale bar =100um. DAPI-grey, ACE2 (magenta), SFTPC (teal), & CD31-Human (green).

**Figure S2. Related to Figure 1. Inoculation of NRG-L fLX mice with SARS-CoV-2.**

**A.** Gating strategy for analyzing the human hematopoietic compartment in naïve fLX following engraftment.

**B.** Representative flow cytometry dot plots delineating the basic architecture of human hematopoietic compartment in fLX (top, lung xenograft) and spleen (bottom) of non-infected NRG-L mice.

**C.** Frequencies of human and mouse CD45^+^ cells in the fLX (top) and spleen (bottom) of non-infected NRG-L mice. n=3 fLX. Mean±SEM.

**D.** Frequencies of human CD3^+^, CD19^+^, CD33^+^ CD11c^+^, and CD56^+^ cells in fLX (top) and spleen (bottom) of non-infected NRG-L mice. n=3 fLX. Mean±SEM.

**E-G.** Probability of survival (E), weight loss (F), and temperature change (G) of NRG-L mice following subcutaneous inoculation with 10^4^ PFU (n=6, blue) or 10^6^ PFU (n=9,red) of SARS-CoV-2, or with PBS (n=1, black).

**H.** Macroscopic representative evidence of pathology in inoculated fLX. (left) Uninoculated control, soft doughy, homogenous pale tan to white fLX. (right) 7 days post SARS-CoV-2 infection, areas of red (black arrow) and pale tan to yellow (asterisk) were histologically confirmed to represent hemorrhage and coagulative necrosis, respectively.

**I-L.** Representative SARS-CoV-2 N IHC on inoculated fLX tissue section (2DPI: I,J; 7DPI: K,L) from NRG-L mice using 10^4^ PFU of SARS-CoV-2. I,K: 200x, scale bar=100µm; J,L: 400x, scale bar=50 µm.

**M-O.** Representative SARS-CoV-2 N IHC on naive (M) and inoculated fLX tissue section (2DPI: N; 7DPI: O) from NRG-L mice using 10^6^ PFU of SARS-CoV-2. 400x, scale bar=50 µm.

**P.** Representative SARS-CoV-2 N IHC on inoculated fLX tissue section (2DPI) from NRG-L mice using 10^6^ PFU of SARS-CoV-2, with a focus on the bronchiole epithelium. 200x, scale bar=100 µm.

**Q.** Representative SARS-CoV-2 N IHC on non-inoculated contralateral fLX tissue section (7DPI) from infected NRG-L mice using 10^6^ PFU of SARS-CoV-2. 200x, scale bar=100 µm.

**R.** SARS-CoV-2 viral RNA quantification by RT-qPCR in the serum of non-infected (naïve) and infected animals at 2DPI and 7DPI using 10^6^ SARS-CoV-2 PFU (n=4-10 fLX). Limit of detection (LOD) is shown as a dotted line and represent mean viremia (n=4) in non-infected mice. n=4-10. Mean±SEM, Kruskal-Wallis test, ns, non-significant.

**S-V.** Planar *in vivo* imaging of SARS-CoV-2 infection in an NRG-L mouse, following inoculation with rSARS-CoV-2-NL (10^6^ PFU) in the left fLX. These images were used to calculate bioluminescent signals (photons/second/mm^3^) reported in Figure 1S-T, using defined regions of interest (ROI) that are shown in panel S.

**Figure S3. Related to Figure 2. Histopathological characterization of SARS-CoV-2 infection.**

**A.** Histopathologic score of ten specific histopathological manifestations observed in fLX inoculated with 10^4^ or 10^6^ PFU of SARS-CoV-2 at 2DPI and 7DPI. Sum of the ten score for each mouse was used to calculate cumulative histopathologic score shown in Figure 2A. n=5-10 fLX. mean±SEM, two-way ANOVA \**p*≤0.05, \*\**p*≤0.01.

**B.** Histopathologic score of ten specific histopathological manifestations observed in fLX inoculated with 10^6^ PFU of SARS-CoV-2 at 2DPI and 7DPI in comparison to naïve/Contralateral (CL) fLX. Sum of the ten score for each mouse was used to calculate cumulative histopathologic score shown in Figure 2B. n=8-12 fLX. mean±SEM, Kruskal-Wallis test \**p*≤0.05, \*\**p*≤0.01, \*\*\**p*≤0.001.

**C-N.** Temporal analysis of SARS-CoV-2 infection in the lungs of K18-hACE2 mice prior infection (C-E), at 2DPI (F-H,L) and 7DPI (I-K,M-N). Representative (n=3 mice/group) histological changes (D,G,J) and SARS-CoV-2 Spike antigen (E,H,K,L-N) distribution and abundance are shown. (C-E) PBS/Sham uninfected controls had no appreciable histologic changes and SARS-CoV-2 S antigen was not observed. (F-H, L) At 2DPI, mild to moderate multifocal interstitial pneumonia with occasional reactive endothelium lining blood vessels and abundant SARS-CoV-2 S antigen within alveolar type 1 & 2 pneumocytes (AT1 & AT2). (I-K,M-N) At 7DPI, increased interstitial histiocyte and lymphocyte infiltrates with lesser numbers of neutrophils. SARS-CoV-2 Spike was observed in histologically normal lung tissues but not in areas of prominent inflammation. H&E and DAB (SARS-CoV-2 Spike antigen), 50X (C,F,I: scale bar = 400 μm), 100X (D,G,J,E,H,K: scale bar = 200 μm), 200x (L,M: scale bar = 100 μm), 400x (N: scale bar = 50 μm) total magnification.

**Figure S4. Related to Figure 2. Ultrastructural characterization of SARS-CoV-2 infection.**

**A.** Suspected viral budding events (asterisks). Inset magnification is shown at the top right. Spike proteins can be observed (arrow).

**B.** Binucleate type II pneumocyte syncytial cell with chromatin condensation and nuclear pyknosis with extracellular virions admixed with necrotic cellular debris and lamellar bodies.

**C-D.** (C) Dying pneumocyte in the airspace filled with viral particles being released in the extracellular milieu. (D) Magnified inset (2X) from panel C.

**E.** Virus replication centers within the cytoplasm of type II pneumocyte.

**F.** Occlusion of vascular lumen by polymerized fibrin. Adjacent endothelium is abruptly absent suggestive of necrosis.

**Figure S5. Related to Figure 3. Transcriptomic and proteomic characterization of SARS-CoV-2 infection in NRG-L mice.**

**A-B.** Host transcript frequencies (A; human, mouse, and SARS-CoV-2) and SARS-CoV-2 gene counts (B) in naïve fLX (n=4), as well as in 2DPI (n=4), 7DPI (n=8) and contralateral fLX (n=3) used for bulk transcriptomic.

**C.** PCA plots of 2DPI (left; n=4), 7DPI (middle; n=6) and contralateral (right; n=3) samples vs. naïve samples following bulk transcriptomic analysis.

**D.** Number of downregulated (left) and up-regulated (right) transcripts overlapping or not between 2DPI (n=4), 7DPI (n=6) and contralateral (n=3) fLX samples.

**E.** Normalized count of IFNB1 and IFNL1 transcripts in naïve, 2DPI (n=4), 7DPI (n=6) and contralateral (n=3) fLX samples. Adjusted p-values in comparison to naïve fLX are indicated.

**F.** Cluster heatmap representing protein up-(z-score>0) and downregulated (z-Score<0) in NRG-L fLX at 7DPI (n=7, 10^6^ PFU of SARS-CoV-2) in comparison to naïve fLX (n=5).

**G.** Cluster heatmap representing protein up-(z-score>0) and downregulated (z-Score<0) in NRG-L fLX at 7DPI (n=7, 10^6^ PFU of SARS-CoV-2) in comparison to 2DPI (n=4) fLX.

**H.** Protein pathway enrichment analysis at 7DPI in comparison to naïve fLX. Level of enrichment for each protein pathway is colored coded.

**Figure S6. Related to Figure 3. Phospho-proteomic characterization of SARS-CoV-2 infection in NRG-L mice.**

**A-B.** Cluster heatmap representing proteins with up- (z-score>0) and down-phosphorylation (z-Score<0) in NRG-L fLX at 2DPI (A; n=4) and at 7DPI (B; n=7) (10^6^ PFU of SARS-CoV-2) in comparison to naïve (n=5) fLX.

**C-D.** Volcano plots displaying differentially phosphorylated proteins at 2DPI (C, n=4) and 7DPI (D, n=7) in inoculated fLX of NRG-L mice in comparison to naïve (n=5) fLX. Proteins with *p*≤0.05 (horizontal dashed line) and with logFC≥1 or ≤-1 (vertical dashed lines) are considered significantly up- or downregulated respectively.

**E-F.** Protein pathway enrichment analysis at 2DPI (E; n=4) and 7DPI (F; n=7) in comparison to naïve fLX (n=3). Level of enrichment for each protein pathway is colored coded.

**Figure S7. Related to Figure 5 and 6. Characterization of SARS-CoV-2 infection in HNFL mice.**

**A.** t-SNE plots displaying clustered expression (scaled expression) of several transcripts coding for several human myeloid, inflammatory, and regulatory markers (CD33, CD14, CD68, IL1β, IL10) in naïve HNFL fLX (n=2). Cluster defining genes (i.e. whose expression level is significantly associated with a given cluster) have their name followed with an asterisk, and Log2FC value are indicated near the corresponding cluster(s). MP, macrophages.

**B.** Volcano plots displaying differentially expressed proteins in naïve HNFL (n=4) vs. NRG-L (n=4) fLX following side-by-side analysis by mass spectrometry run. Proteins with *p*≤0.05 (horizontal dashed line) and with logFC≥1 or ≤-1 (vertical dashed lines) are considered significantly up- or downregulated respectively.

**C.** Quantification of tissue area immunoreactive for CD68 (% of analyzed tissue) using Halo Image analysis in naïve fLX from HNFL and NRG-L mice. n=3-8. Mean±SD, Welch’s t test \*\*\*\**p*≤0.0001.

**D.** SARS-CoV-2 viral RNA quantification by RT-qPCR in inoculated fLX of NRG-L or NRGF-L (no HSC engraftment) at 2DPI following SARS-CoV-2 inoculation (10^6^ PFU) (n=3 fLX per group). Limit of detection is showed as a dotted line (LOD) and represent mean viral load (n=7) in naïve fLX extracted from NRG-L and NRGF-L mice. Mean±SEM, Kruskal-Wallis test, ns non-significant.

**E.** Histopathologic score of four selected specific histopathological manifestations observed in NRG-L (n=8-12) and HNFL (n=4) fLX inoculated with 10^6^ PFU of SARS-CoV-2 at 2DPI and 7DPI. Sum of all the histopathologic score recorded for each fLX analyzed was used to calculate cumulative histopathologic score shown in Figure 5K. Mean±SEM, two-way ANOVA \**p*≤0.05, \*\**p*≤0.01, \*\*\**p*≤0.001.

**F.** Gating strategy (Flow cytometry dot plots) used to delineate the human hematopoietic compartment in naïve and inoculated HNFL fLX.

**G-H.** Volcano plots displaying differentially phosphorylated proteins at 2DPI in inoculated fLX of NRG-L mice (F) and HNFL mice (G) following side-by-side sample analysis by mass spectrometry. Proteins with *p*≤0.05 (horizontal dashed line) and with logFC≥1 or ≤-1 (vertical dashed lines) are considered significantly up- or downregulated respectively. Naïve, n=4; 2DPI, n=4.

**I.** Significantly (*p*≤0.05) differentially expressed transcripts (upregulated, red; downregulated, blue) in inoculated HNFL fLX at 7DPI (n=3) following SARS-CoV-2 infection (10^6^ PFU) in comparison to naïve HNFL fLX (n=2). Fold changes were derived from single-cell RNAseq datasets and were computed using MAST. Transcripts with *p*≤10^-100^ (horizontal dotted line) and with log_2_ fold change≥0.2 or ≤-0.2 (vertical dotted lines) are highlighted.

**J-K.** t-SNE plots displaying clustered expression (scaled expression) of several transcripts coding for human cell surface markers, cytokines and ISGs in the human compartment of HNFL fLX at 7DPI (J) and at 2DPI (K). (J) Myeloid, inflammatory, and regulatory markers expression (CD33, CD14, CD68, IL1β, IL10) in inoculated fLX at 7DPI. (K) ISG expression (OAS1-2, MX2, DDX58 and IFI44L) in inoculated HNFL fLX at 2DPI. Cluster of interest are indicated with a dotted circle, and each cluster name is indicated at the left of each panel (AIM, activated inflammatory macrophages; ARM, activated regulatory macrophages; SSM, steady-state macrophages; DC, dendritic cells; Fib, fibroblasts; Mon, monocytes). Cluster defining genes (i.e. whose expression level is significantly associated with a given cluster) have their name followed with an asterisk, and Log2FC value are indicated near the corresponding cluster(s). Naïve, n=2; 2DPI, n=3; 7DPI=3.

**L.** Viral gene count per cluster, segregated between activated macrophage clusters (AM) and all other (Others). n=29 clusters. Mean±SD, Mann-Whitney t-test, \*\*\**p*≤0.001.

### Supplemental items

**Supplemental item 1. Related to Figure 1,4 and 6.** Single-cell RNA sequencing gene-defining clusters and cluster annotation (Excel file).

**Supplemental item 2. Related to Figure 1.** Time-lapse (2DPI, 4DPI, 6DPI and 12DPI) 3D *in vivo* imaging of SARS-CoV-2 infection (mp4 file)

**Supplemental item 3. Related to Figure 3.** List of differentially expressed genes and IPA scores from bulk RNA sequencing analysis (Excel file)

**Supplemental item 4. Related to Figure 3.** Proteomic analysis Matrix_NRG-L only (Excel file) **Supplemental item 5. Related to Figure 3.** Phospho-proteomics analysis Matrix_ NRG-L only (Excel file)

**Supplemental item 6. Related to Figure 5.** Proteomic analysis Matrix_NRG-L vs. HNFL (Excel file)

**Supplemental item 7. Related to Figure 5.** Phospho-proteomics analysis Matrix_NRG-L vs. HNFL (Excel file)

