## Supplementary figures and images for "Macrophages govern antiviral responses in human lung tissues protected from SARS-CoV-2 infection"

### Supplemental Figure 1

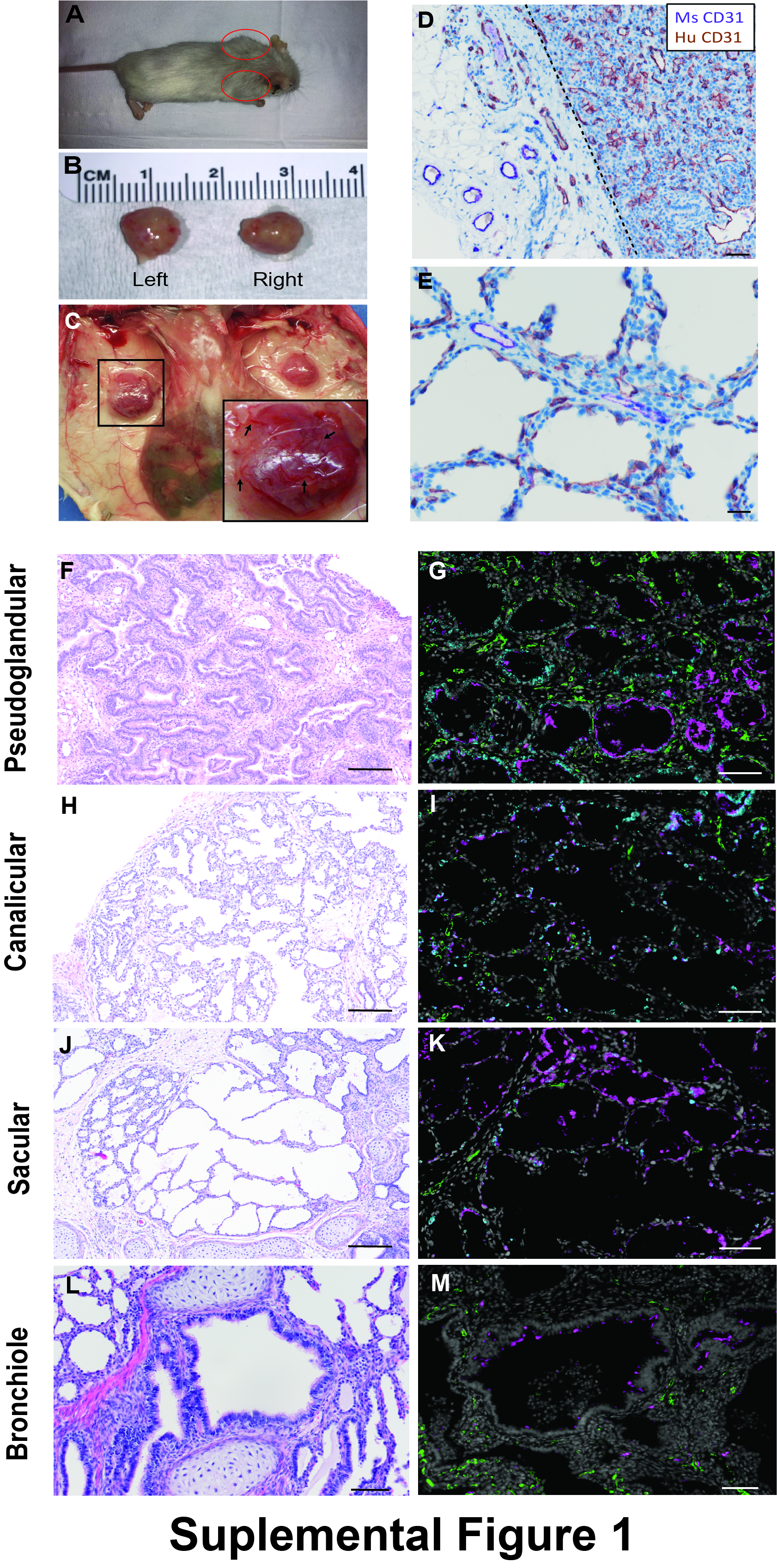

### Supplemental Figure 2

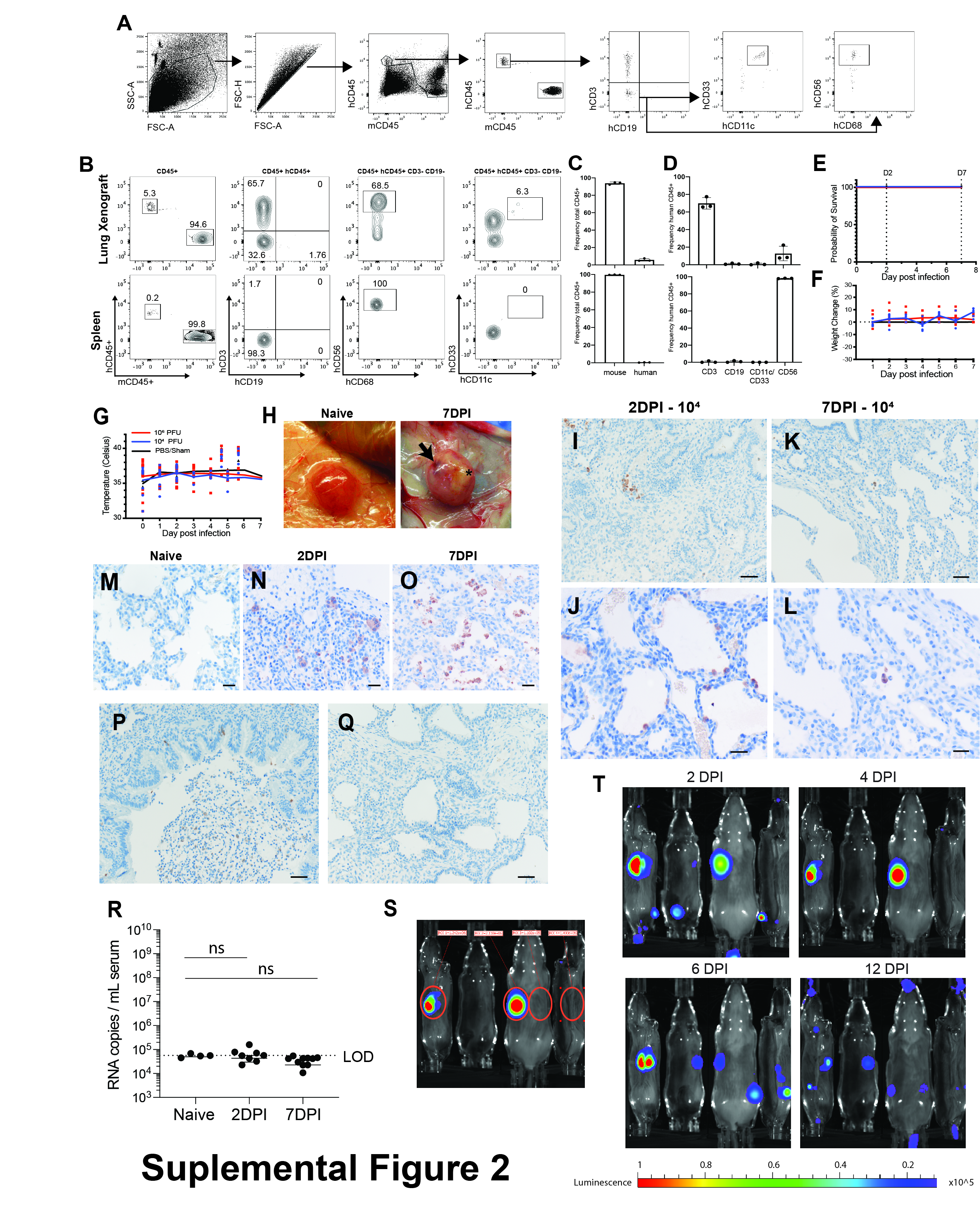

### Supplemental Figure 3

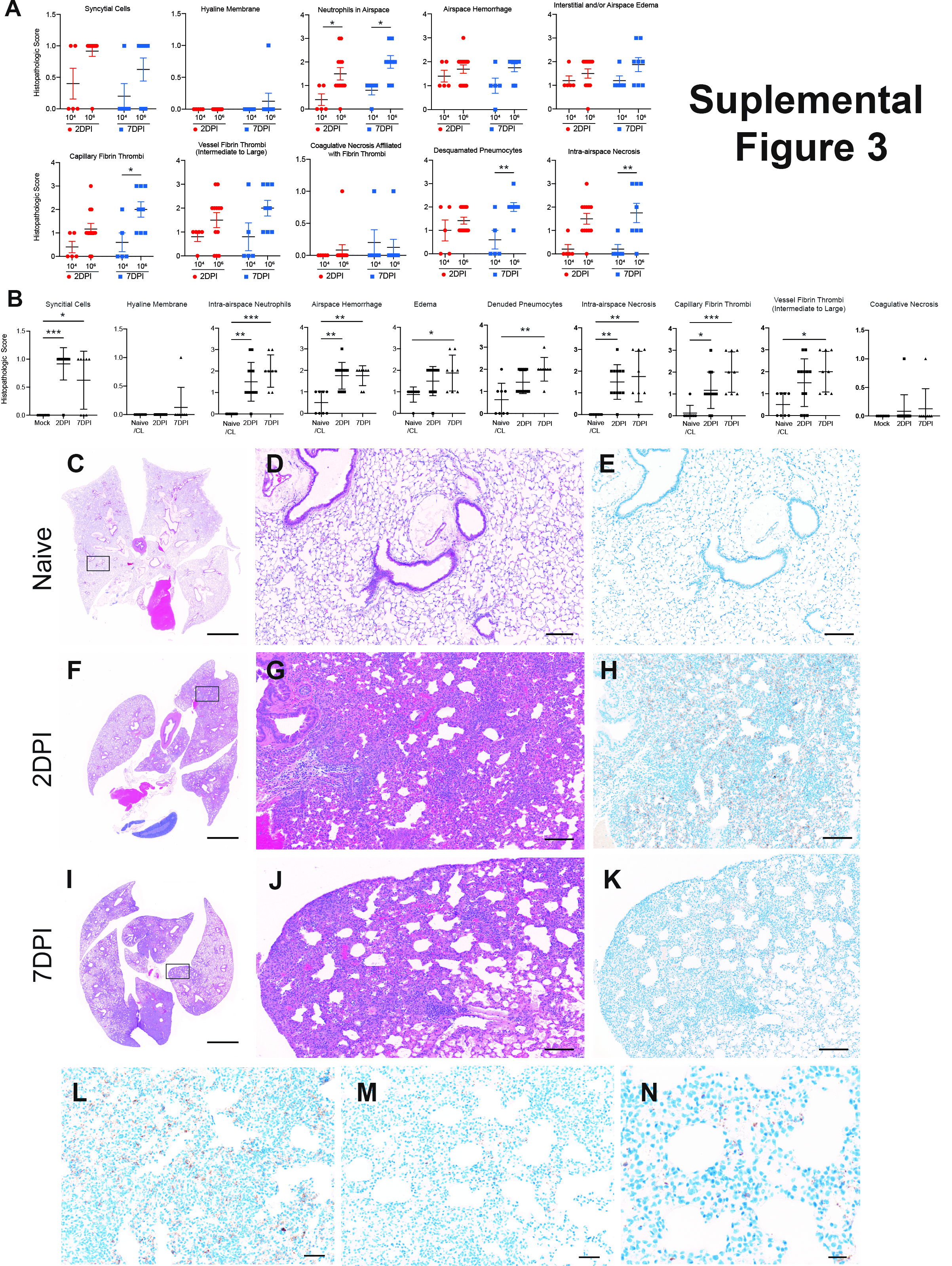

### Supplemental Figure 4

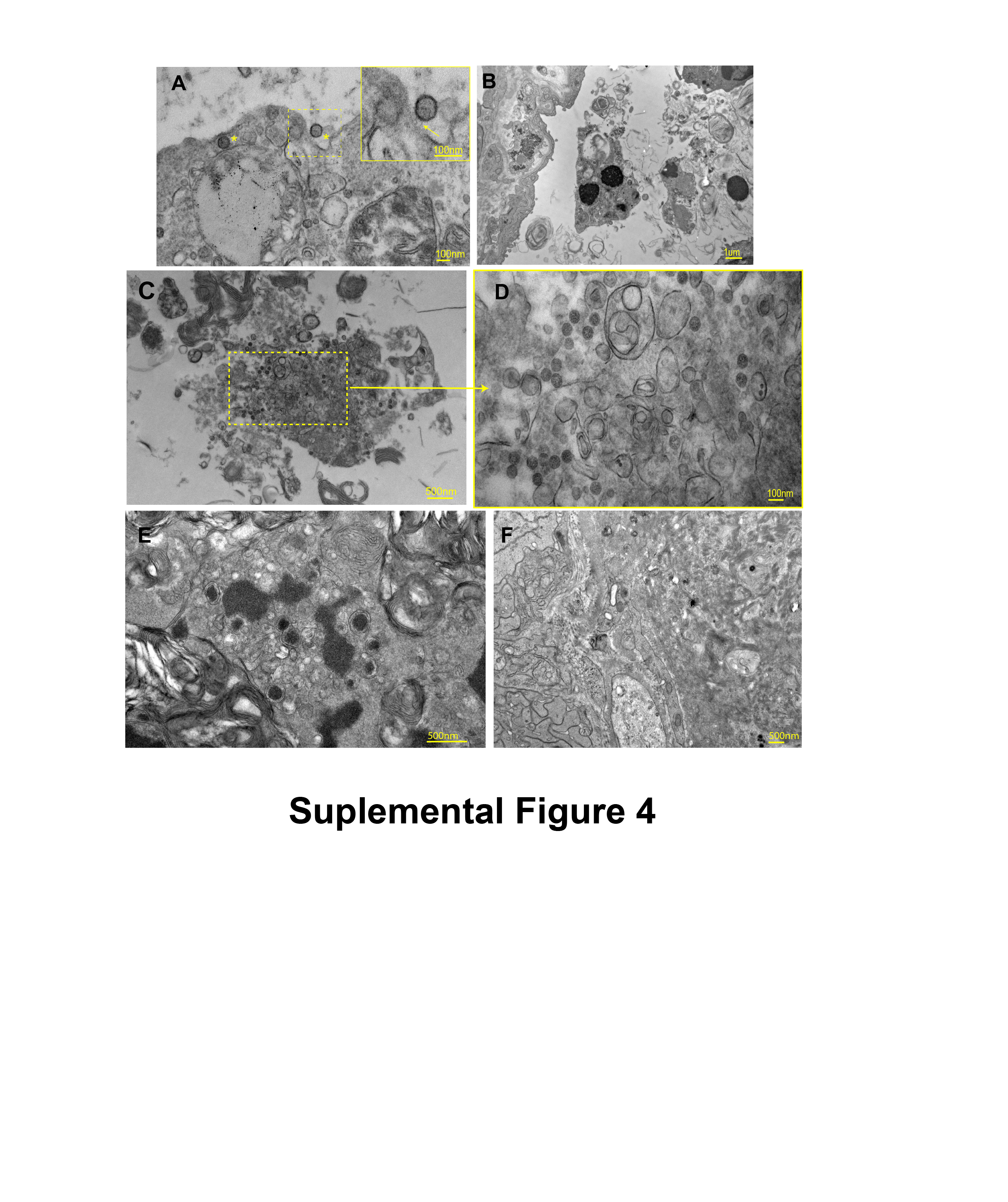

### Supplemental Figure 5

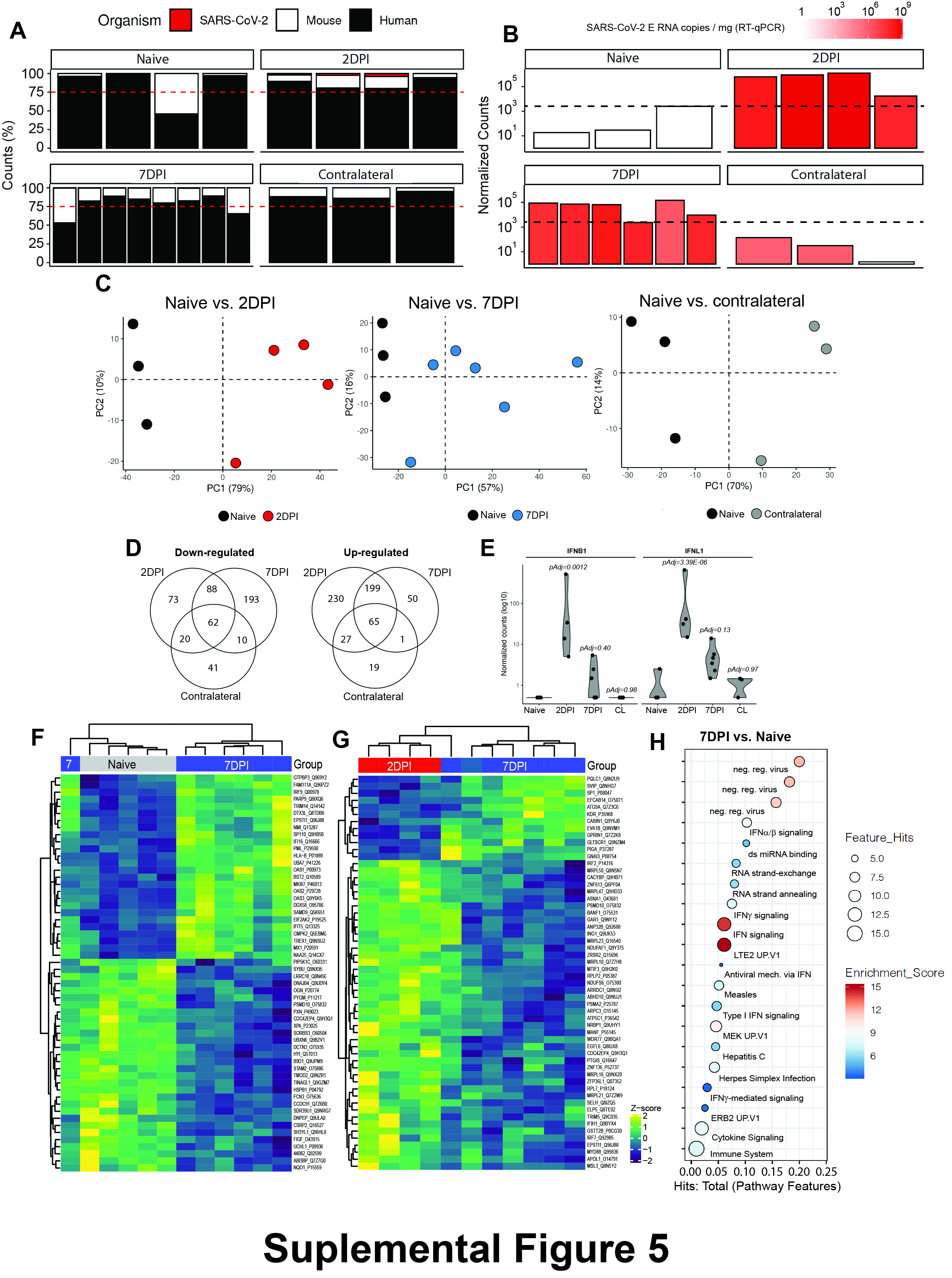

### Supplemental Figure 6

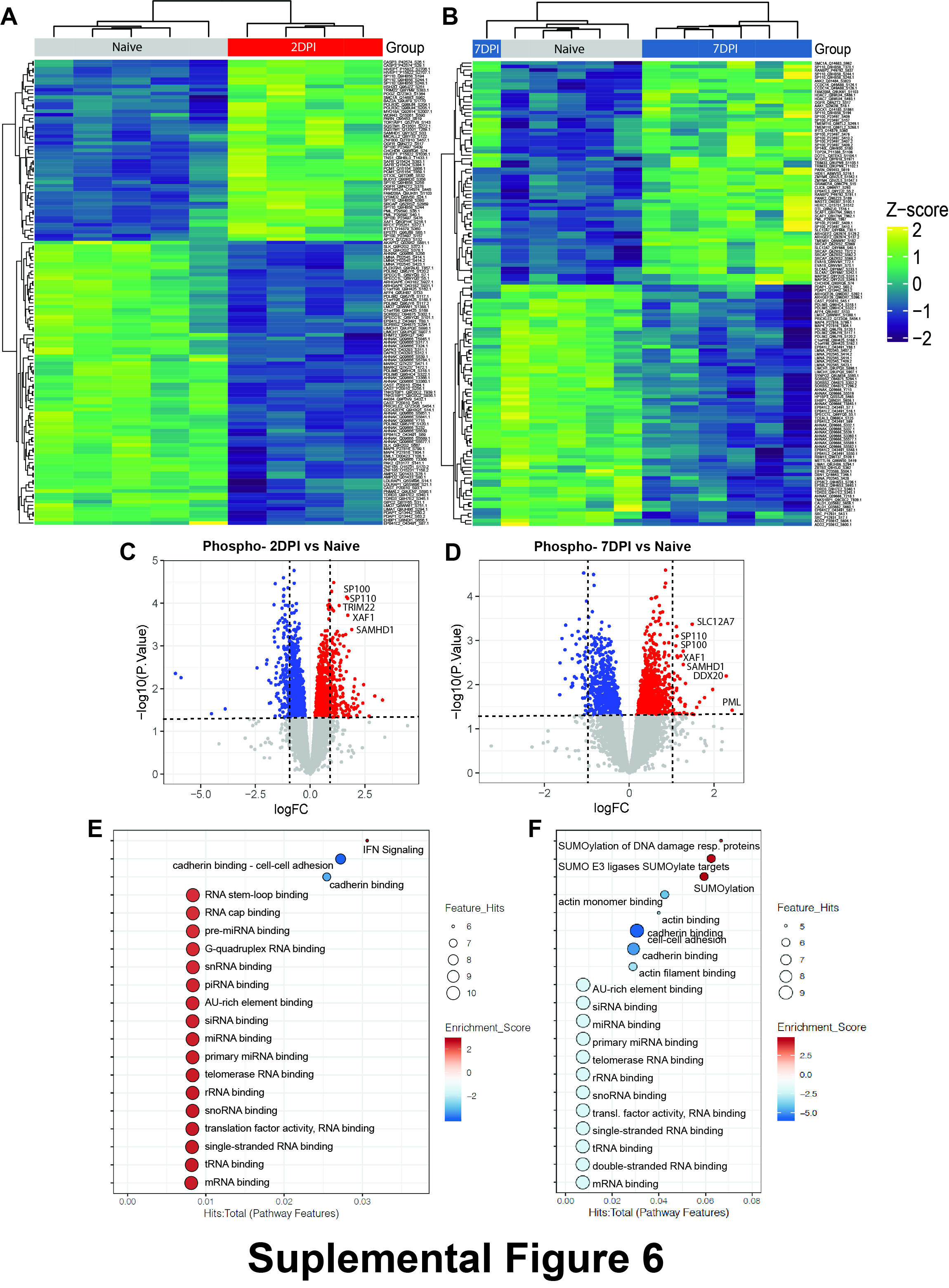

### Supplemental Figure 7

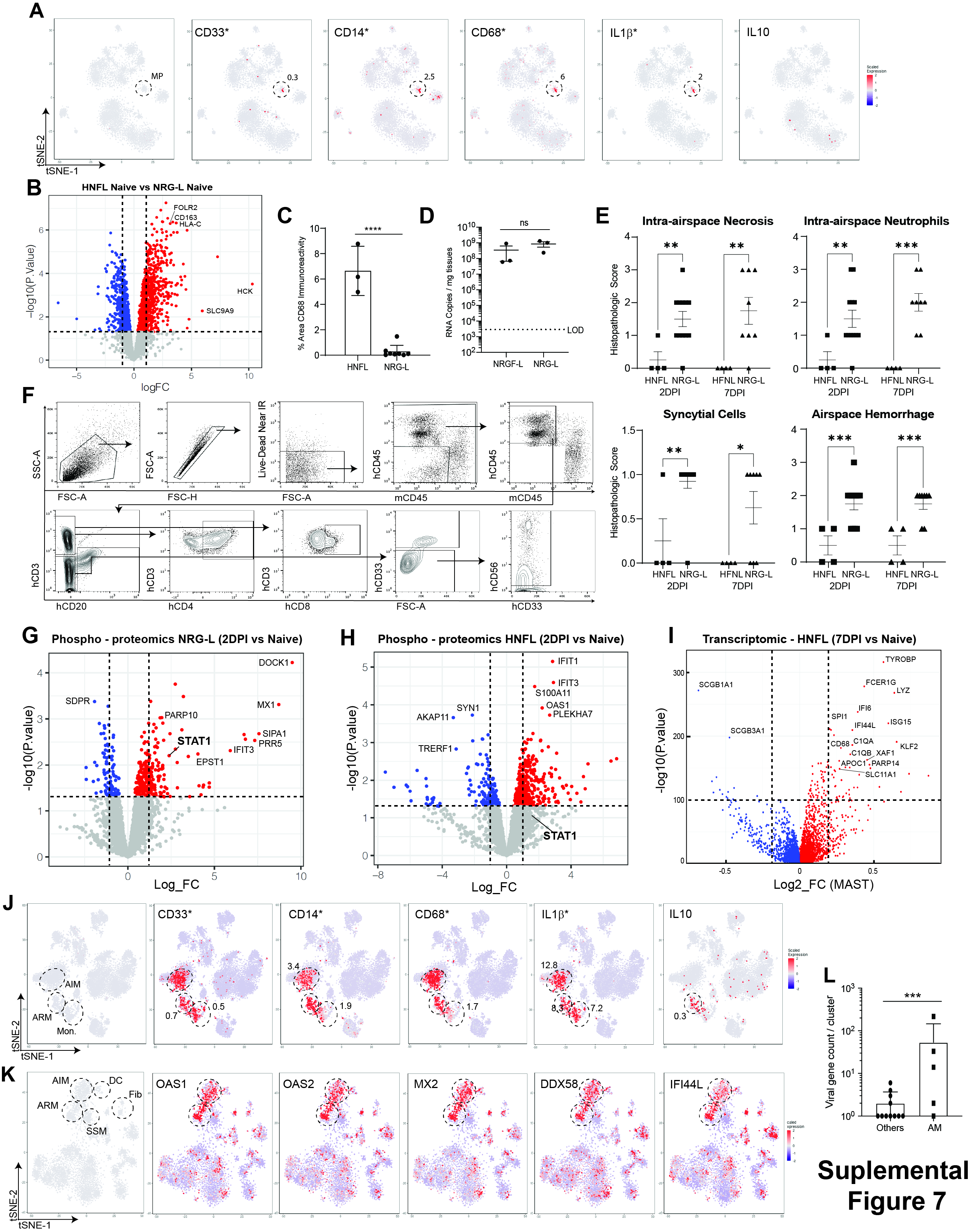
